# DuoHexaBody-CD37 induces direct cytotoxic signaling in diffuse large B-cell lymphoma

**DOI:** 10.1101/2025.02.24.639899

**Authors:** Simar Pal Singh, Kumar Mangalam, Michelle D. van den Beukel, Sjoerd van Deventer, Marije B. Overdijk, M. Guy Roukens, Kim C.M. Santegoets, Esther C.W. Breij, Martin ter Beest, Willem P. J. Cox, Annemiek B. van Spriel

**Author notes:** Corresponding author: Annemiek van Spriel. These authors contributed equally (Kumar Mangalam, Michelle D. van den Beukel).

## Abstract

Diffuse large B-cell lymphoma (DLBCL) is a common aggressive form of Non-Hodgkin lymphoma. Tetraspanin CD37 is highly expressed on mature B cells and being studied as a therapeutic target for NHL, including DLBCL. DuoHexaBody-CD37 is a biparatopic antibody with an E430G hexamerization-enhancing mutation targeting two non-overlapping CD37 epitopes shown to promote complement-dependent cytotoxicity. However, the impact of DuoHexaBody-CD37 on direct cytotoxic signaling has not yet been studied. Here we demonstrate that DuoHexaBody-CD37 induces direct cytotoxicity in DLBCL-derived tumor cell lines independent of the subtype. DuoHexaBody-CD37 induced significant CD37 clustering and was retained at the cell surface in contrast to rituximab, which was internalized. Unbiased screening identified the modulation of 26 (phospho)proteins upon DuoHexaBodyCD37 treatment of primary B cells or DLBCL cells. Whereas DLBCL cells predominantly upregulated p-SHP1(Y564) upon DuoHexaBody-CD37 treatment, primary B cells showed significantly increased p-AKT(S473) and MAPK signaling which is linked to cell survival. Studies using CD37-mutants identified the N-terminus to be involved in DuoHexaBody-CD37-induced signaling. Finally, DuoHexaBody-CD37 treatment inhibited cytokine pro-survival signaling in DLBCL cells. These findings provide novel insights into the signaling functions of CD37 upon DuoHexaBody-CD37 treatment, and open up opportunities for developing CD37-targeted immunotherapy in combination with small molecule inhibitors to maximize tumor cell death.

## Introduction

Diffuse large B-cell lymphoma (DLBCL), an aggressive type of mature B-cell lymphoma, accounts for approximately one third of Non-Hodgkin lymphoma (NHL) cases. DLBCL arises from mature B cells in the lymph node, and comprises a heterogenous group of tumors that can be classified according to cell-of-origin (germinal-center B-cell (GCB) derived and activated B-cell (ABC) subtypes). In addition, recent comprehensive molecular profiling has identified five to seven genetic subtypes that share similar oncogenic pathways (reviewed in ^1^). A large fraction of DLBCL patients (up to 40%) do not respond or relapse after first line treatment with chemotherapy (cyclophosphamide, hydroxydaunorubicin, vincristine sulphate (Oncovin), and prednisone: CHOP) and immunotherapy (rituximab). Rituximab and also next-generation antibodies (ofatumumab) target CD20 at the plasma membrane of DLBCL ^2^, however decreased expression of CD20 is related to inferior clinical outcome after treatment with R-CHOP ^3,4,5^. Thus, targeting alternative B-cell membrane proteins and development of novel immunotherapies are needed to improve clinical outcomes of patients with relapsed/refractory DLBCL ^6,7^.

CD37 is a tetraspanin protein with four-transmembrane domains, which is predominantly expressed by mature B cells ^8–10^. Tetraspanins interact with immune receptors on the same cell (in cis) and control membrane organization in lymphocytes ^11–13^. CD37 is absent on progenitor B cells and terminally differentiated plasma cells making it an ideal target for mature B cell malignancies ^14,15,16^. In DLBCL patients, CD37 was reported to be an independent prognostic factor for both GCB and ABC subtypes ^4^. In line with these findings, CD37-deficient mice show defects in humoral and cellular immune responses, and spontaneously develop mature B-cell lymphoma, which is dependent on IL-6 ^17^. IL-6 signals through the IL-6 receptor complex that activates AKT kinase and STAT3 signaling stimulating cell survival and proliferation. IL-6 signaling has been reported to be a critical driver in the tumor microenvironment and a negative prognostic factor in diffuse large B-cell lymphoma^18^.

CD37 is under preclinical and clinical investigation as novel therapeutic target for BNHL, including antibody-based and chimeric antigen receptor (CAR) T-cell therapies ^19–23,24^. DuoHexaBody-CD37 (GEN3009) is a novel biparatopic immunoglobulin G1 (IgG1) with a point mutation in the Fc-domain (E430G) that enhances antibody hexamerization upon binding to the cell surface ^25^. This facilitates C1q binding and results in potent complement-dependent cytotoxicity (CDC). In addition DuoHexaBody-CD37 can induce Fc-gamma receptor (FcγR) mediated antibody-dependent cellular cytotoxicity (ADCC) and phagocytosis (ADCP) by effector cells. DuoHexaBody-CD37 mediates superior CDC in patient-derived DLBCL cells ex vivo compared to CD20 monoclonal antibodies as shown in preclinical studies ^26^.

In B cells, CD37 has been reported to connect to intracellular signaling pathways via its two intracellular tails ^9,27^. The N-terminal tail of CD37 contains a potential ‘ITIM-like’ motif that may induce SHP1 signaling, whereas the C-terminal tail of CD37 bears a predicted ‘ITAM-like’ motif that may stimulate AKT kinase-dependent survival ^27^. Here we investigated whether DuoHexaBody-CD37 is capable of inducing intracellular signaling that induces cell death of malignant B cells. We report that DuoHexaBody-CD37 mediates direct CD37-mediated signaling and evokes tumor cytotoxicity in DLBCL-derived tumor cells, independent of the presence of complement. DuoHexaBody-CD37 modulated the PI3K-AKT and MAPK signaling pathways, with differences observed between primary B cells and DLBCL tumor cells. Moreover, DuoHexaBody-CD37 inhibited IL-4-dependent p-STAT6 and IL-21-dependent p-STAT3 signaling in DLBCL cells. In conclusion, this study shows a novel mechanism of action of DuoHexaBody-CD37, and provides more insight into CD37 as therapeutic target for B cell malignancies.

## Results

### DuoHexaBody-CD37 mediates direct cytotoxicity in DLBCL cell lines

Induction of apoptosis in B cell leukemia and lymphomas has been reported for therapeutic antibodies such as obinutuzumab ^28^, alemtuzumab ^29^, otlertuzumab ^30^, and rituximab ^31^. To investigate the direct cytotoxic potential of DuoHexaBody-CD37, DLBCL-derived cell lines classified as GCB (Oci-Ly8 and Oci-Ly7), ABC (HBL-1 and U2932) and Burkitt lymphoma-derived cell lines (BJAB, Daudi) were treated with DuoHexaBody-CD37 in the absence of complement, and with or without an Fc-targeting F(ab)2-fragment (a-Fc) to facilitate Fc-mediated crosslinking. Whereas the viability of Burkitt lymphoma-derived cell lines (BJAB and Daudi) was not significantly affected by DuoHexaBody-CD37 and Fc-crosslinking treatment, all tested DLBCL-derived cell lines showed >25% reduction in viability after DuoHexaBody-CD37 treatment with a-Fc (Figure 1A). We confirmed that apoptosis was the mechanism of cell death by Annexin V and 7-AAD staining in Oci-Ly7 and HBL1 cell lines upon Fc-crosslinking of DuoHexaBody-CD37 (Supplementary Figure 1A). Notably, differences in viability loss after treatment did not correlate with differences in CD37 membrane expression in the tested cell lines (Supplementary Figure 1B). To investigate the physiological relevance of Fc-crosslinking mediated DuoHexaBody-CD37 induced killing, cytotoxicity assays of DLBCL cell lines were performed using peripheral blood mononuclear cells (PBMCs) expressing Fcɣ receptors (FcɣR). In these assays, PBMCs were pre-fixed to prevent FcɣR signaling and to avoid any influence of ADCC/ADCP as previously shown with DuoHexaBody-CD37 ^25^. DuoHexaBody-CD37 treatment resulted in significantly increased direct cytotoxicity of DLBCL cells compared to untreated cells in presence of PBMCs as source for Fc-crosslinking for all donors tested (Figure 1B). Collectively, these results show induction of direct cytotoxicity by DuoHexaBody-CD37 treatment with Fc crosslinking in different DLBCL cells. To further investigate the contribution of specific FcγRs in effector cell mediated Fc-crosslinking, total PBMCs were compared with defined immune cell subsets: B cells (expressing FcγRIIb), NK cells (expressing FcγRIIIa and FcγRIIc), monocytes (expressing FcγRI, FcγRIIa, FcγRIIb, and FcγRIIIa), and T cells with no confirmed expression of FcγR. All immune cell subsets expressing FcγRs led to similar or greater direct cytotoxicity of DLBCL cells compared with the total PBMC pool (Figure 1C). These results indicate that DuoHexaBody-CD37-induced killing through Fc-mediated crosslinking is not dependent on the expression of specific FcγR subtypes.

**Figure 1:**
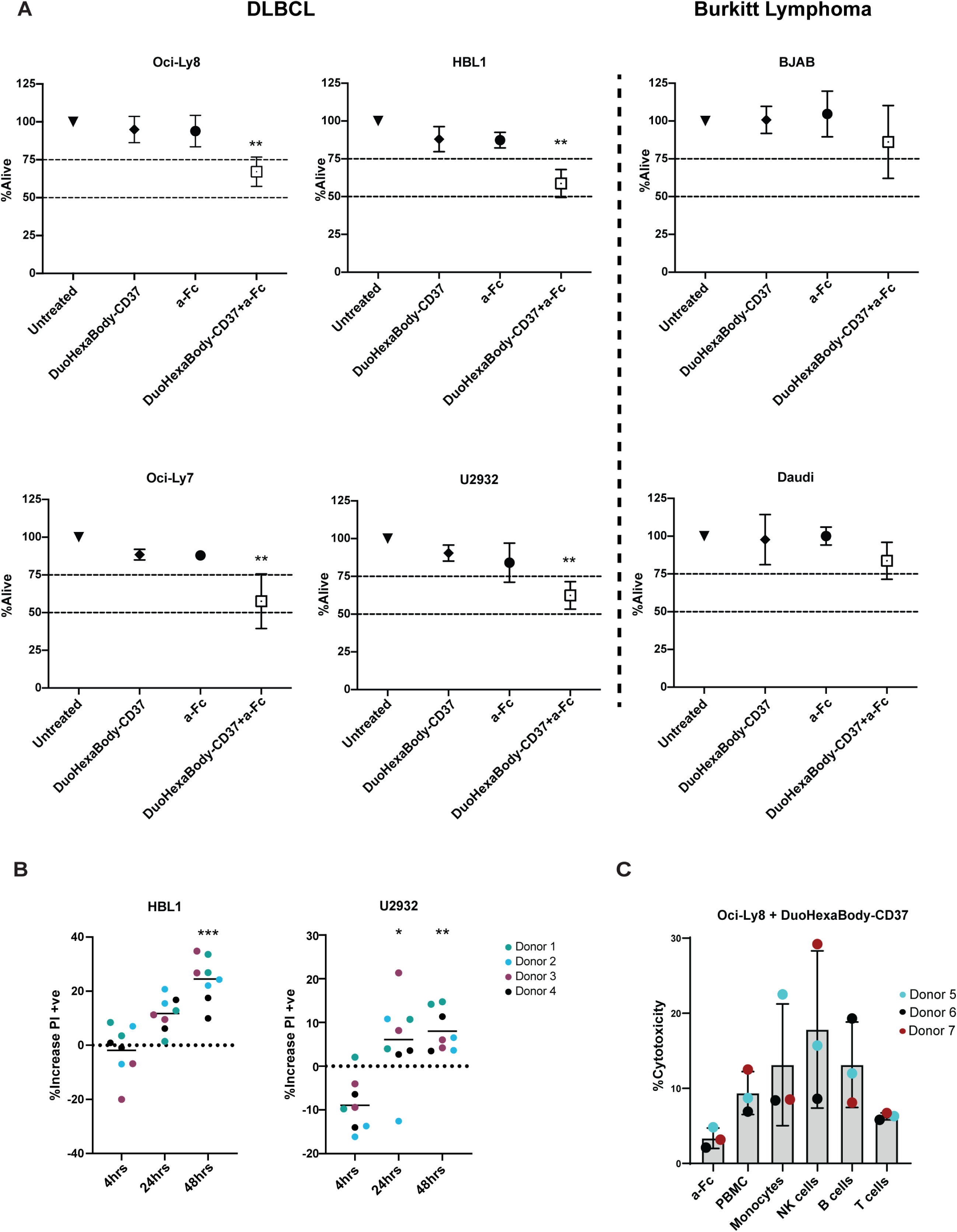
DuoHexaBody-CD37-induces direct cytotoxicity in DLBCL-derived tumor cell lines. **(A)** Percentage decrease (mean +/- SD) in viability upon DuoHexaBody-CD37 treatment in presence or absence of goat F(ab’)2 anti-human IgG (a-Fc) in indicated cell lines after 48h. Data is shown from at least three independent experiments. Significance was calculated compared to untreated control using Kruskal-Wallis test followed by Dunn’ multiple comparison test correction (**p<0.01) **(B)** Percentage increase in cell death (PI-positive) upon co-culturing healthy donor-derived fixed PBMCs with DuoHexaBody-CD37 pre-treated HBL-1 and U2932 cells compared to untreated cells for indicated time points. Duplicates from each individual donor (n=4) are indicated in separate colors. Significance was calculated compared to 4h treated cells using Kruskal-Wallis test followed by Dunn’ multiple comparison test correction (*p<0.05, **p<0.01, ***p<0.001). **(C)** Percentage of cell death in DuoHexaBody-CD37 pre-treated Oci-Ly8 cells (eFluor780-positive) upon co-culturing with healthy donor-derived fixed PBMC, monocytes, NK cells, B cells and T cells compared to Fc-crosslinker. Measurements from three individual donors (mean +/- SD) are shown, with each donor represented by a distinct colored symbol.

### DuoHexaBody-CD37 induces CD37 clustering without modulating CD37 cell surface expression

CD37 is involved in the spatial organization of the B-cell plasma membrane by forming tetraspanin nanodomains ^13^, therefore we investigated CD37 membrane clustering on DuoHexaBody-CD37-treated tumor cells compared to tumor cells treated with the IgG1 isotype control antibody with and without a-Fc mediated crosslinking, respectively. Whereas CD37 surface expression was the same in both samples, the clustering of CD37 on tumor cells (measured by fluorescence intensity/area) was significantly higher upon DuoHexaBody-CD37 treatment than isotype control antibody treatment (Figure 2A, B). Interestingly, DuoHexaBody-CD37-mediated clustering of CD37 was observed in both Burkitt cells (BJAB) (Figure 2A) and DLBCL (Oci-Ly8) (Figure 2B) even in absence of Fc-crosslinker. Upon crosslinking, clustering of CD37 increased in both cell lines, however the effect was more pronounced in BJAB cells compared to Oci-Ly8. Thus, DuoHexaBody-CD37 induces potent CD37 membrane clustering at the cell surface of malignant B cells with and without Fc-mediated crosslinking.

**Figure 2:**
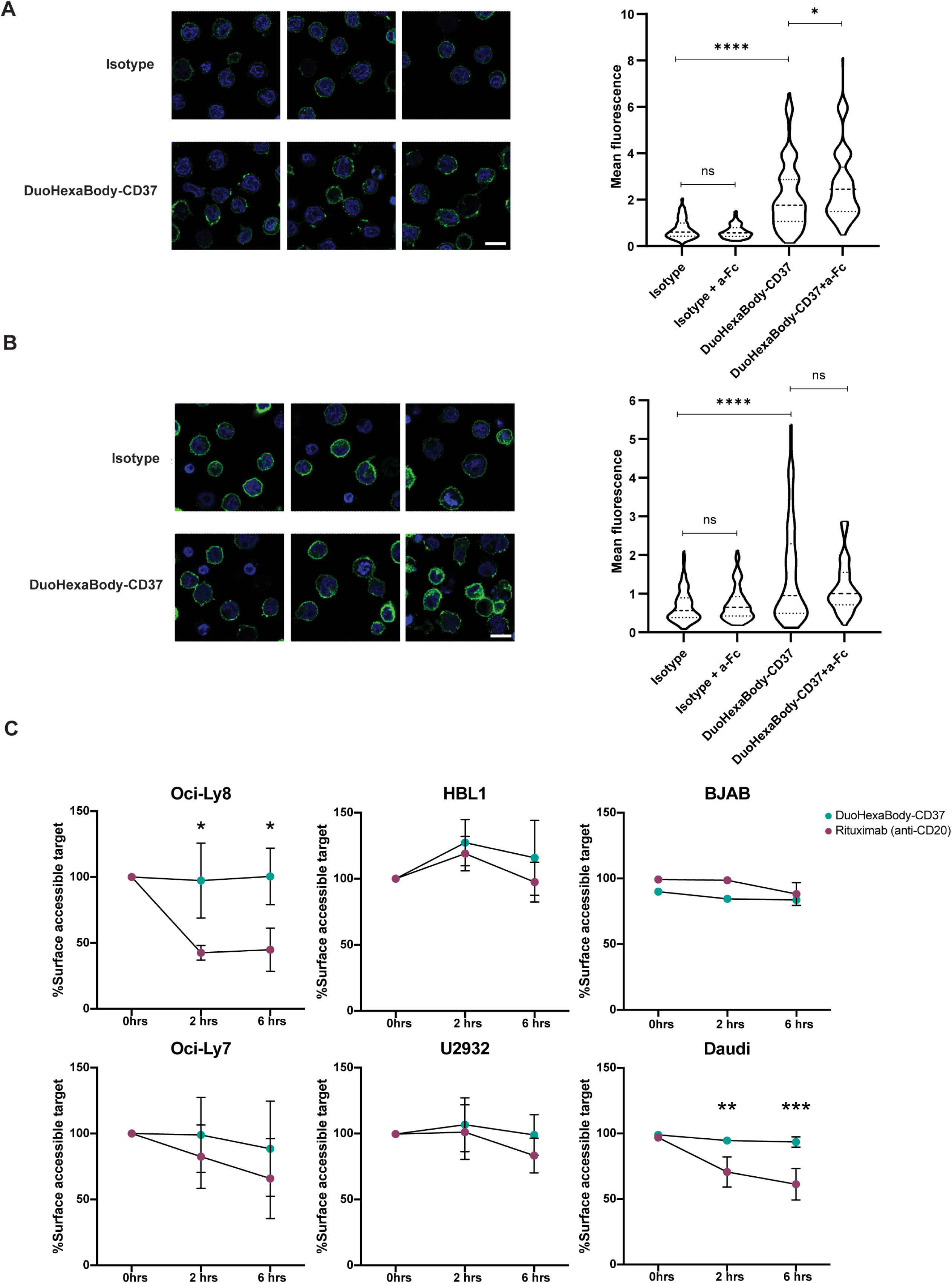
DuoHexaBody-CD37 induces CD37 clustering without modulating CD37 cell surface expression. **(A,B)** Airyscan images depicting clustering (fluorescence/area) upon DuoHexaBody-CD37 (DHB-CD37) treatment or B12 isotype control in BJAB **(A)** and Oci-Ly8 **(B)** cells with or without Fc-crosslinking (a-Fc). Data shown from five independent experiments. Bar is 10 µm. Significance was calculated comparing DuoHexaBody-CD37-treated cells to isotype control antibody in the presence and absence of Fc-crosslinking (a-Fc) respectively, using Kruskal-Wallis test followed by Dunn’s multiple comparisons test (****p<0.0001, *p<0.05). **(C)** Percentage decrease (mean +/- SD) in cell surface binding by DuoHexaBody-CD37 or rituximab (anti-CD20) in indicated cell lines as measured by flow cytometry. Data is shown from at least three independent experiments. Significance was calculated comparing respective time points using unpaired T-test (*p<0.05, **p<0.01, ***p<0.001).

Since the efficacy of rituximab can be negatively impacted by induction of CD20 internalization, leading to loss of CD20 expression at the cell surface ^32,33^, we investigated whether DuoHexaBody-CD37 affected CD37 cell surface accessibility. We observed no changes in surface CD37 expression upon DuoHexaBody-CD37 treatment in all cell lines tested, in contrast to CD20 expression upon rituximab treatment, which was decreased on DAUDI, Oci-Ly8 and Oci-Ly7 cells (Figure 2C). We next assessed CD37 surface availability with and without Fc-crosslinking of DuoHexaBody-CD37 and observed no consistent reduction in CD37 surface availability upon crosslinking (Supplementary Figure 2). Thus, the efficacy of DuoHexaBody-CD37 is not affected by loss of CD37 through internalization of surface molecules.

### DuoHexaBody-CD37 treatment results in activation of multiple downstream signaling pathways

Next, we focused on investigating mechanisms underlying the direct cytotoxicity upon DuoHexaBody-CD37 treatment by analyzing DuoHexaBody-CD37-induced downstream signaling using high throughput Reverse Phase Protein Array (RPPA) ^34^. This technique enables simultaneous unbiased measurement of 484 proteins (102 phospho targets and 382 total proteins) in multiple samples at once. For this analysis primary B cells (purified CD19+ B cells from PBMCs) and two DLBCL-derived cell lines (Oci-Ly7 and U2932) were treated with DuoHexaBody-CD37 in the presence or absence of Fc-crosslinker and compared to untreated samples from each cell type. Principal component analysis on all tested proteins in RPPA identified 3 separate clusters correlating with the different cell models (Supplementary Figure 3). Compared to respective untreated controls, 26 phosphoproteins showed at least a two-fold change in either of the three cell models tested upon DuoHexaBody-CD37 treatment (Figure 3A). Pathway analysis on all 26 phospho-sites identified PI3K/AKT/MTOR signaling as the most significantly enriched pathway upon DuoHexaBody-CD37 treatment (Figure 3B). In addition, some of these signaling proteins are downstream of B cell receptor (BCR) signaling and RAS signaling pathways, which have been reported to be essential for B cell survival and lymphomagenesis ^35,36^.

**Figure 3:**
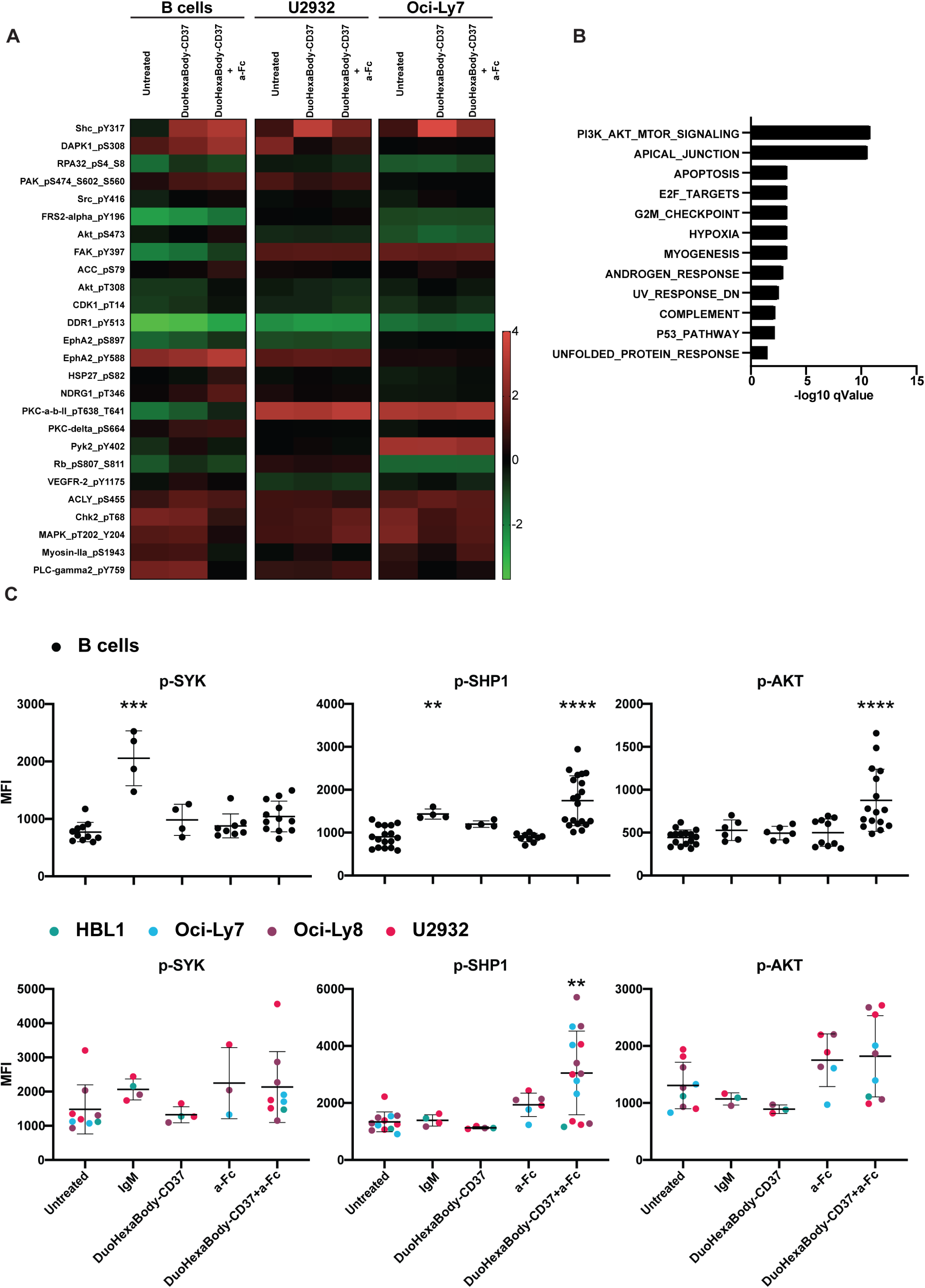
DuoHexaBody-CD37 treatment results in activation of different downstream signaling pathways. (**A**) Heatmap depicting log-normalized signal intensities of 26 phosphoproteins with two-fold increase in signal upon DuoHexaBody-CD37 treatment in presence or absence of goat F(ab’)2 anti-human IgG (a-Fc) in primary B cells, U2932 and Oci-Ly7. (**B**) Oncogenic hallmark signatures obtained from the MSigDB enriched using 26 phosphoproteins with two-fold increase in signal upon DuoHexaBody-CD37 treatment in presence or absence of goat F(ab’)2 anti-human IgG (a-Fc) in primary B cells, U2932 and Oci-Ly7. (**C**) Phosphoflow analysis of pSYK(Y348), p-SHP1(Y564), p-AKT(S473) on primary B cells (*top row*) or DLBCL cells (*bottom row*) either untreated or treated with anti-BCR (F(ab’)2 anti-IgM), DuoHexaBody-CD37 and/or goat F(ab’)2 anti-human IgG (a-Fc). Dot plots depict quantification (mean +/- SD) of mean fluorescence intensity (MFI). Each dot represents individual donor (B cells) or experimental replicate (DLBCL). Significance was calculated compared to untreated control using Kruskal-Wallis test followed by Dunn’ multiple comparison test correction (**p<0.01, ***p<0.001, ****p<0.0001).

Next, the effect of DuoHexaBody-CD37 treatment on primary B cells versus tumor cells was studied by focusing on those phosphoproteins that showed approximately two-fold upregulation in signal compared to untreated controls. Primary B cells showed upregulation of multiple phosphoproteins upon DuoHexaBody-CD37 treatment including p-AKT, p-FAK, pSrc and p-Shc. In DLBCL cells, different phosphoproteins (p-RPA32, p-DAPK1, p-PAK, p-PI3K and p-FRS-alpha) were upregulated upon DuoHexaBody-CD37 treatment. The list of targets and fold changes can be found in Supplementary Table 1.

To validate these results, we analyzed different targets within the three main signaling pathways that came out of the RPPA analysis: the PI3K/AKT pathway, BCR pathway and MAPK/ERK pathway using phosphoflow analysis. Anti-BCR (IgM) was used as stimulation control in the phosphoflow studies. We confirmed specific upregulation of p-AKT(S473) in primary B cells upon treatment with DuoHexaBody-CD37 in the presence of Fc-crosslinker compared to untreated cells (Figure 3C). In DLBCL cell lines, upregulation of p-AKT(S473) was not specific for DuoHexaBody-CD37 treatment as this was already observed in presence of the Fc-crosslinker only (Figure 3C). These data indicate a differential role of DuoHexaBody-CD37 mediated signaling in primary B cells versus DLBCL which could be partially explained by constitutive activation of the PI3K-AKT pathway in DLBCL ^37^. As engagement of cell death pathways in CLL cells was shown to be dependent on SHP1 recruitment and activation via CD37 N-terminus ^27^, we additionally assessed p-SHP1(Y564) status upon DuoHexaBody-CD37 treatment of primary B cells and DLBCL cells via phosphoflow. Compared to untreated cells, DuoHexaBody-CD37 treatment in the presence of Fc-crosslinker induced significant upregulation of p-SHP1(Y564) in both primary B cells and DLBCL cells (Figure 3C, Supplementary Figure 4A).

To analyze phosphoproteins downstream of the BCR, we studied p-SYK, p-BTK, pPLCγ2 and p-PKC in primary B cells and DLBCL cells by phosphoflow. p-SYK was observed to be upregulated only after BCR stimulation (anti-IgM), but not after DuoHexaBody-CD37 treatment of primary B cells or DLBCL cells (Figure 3C). On other hand, p-PLCy2(Y759) and p-BTK(Y223) were specifically upregulated in DLBCL cells upon treatment with DuoHexaBody-CD37 in the presence of Fc-crosslinker (Figure 4A, Supplementary Figure 4B). Next, we analyzed the MAPK signaling pathway and observed p-P38 to be specifically upregulated in primary B cells, but not in DLBCL cells, upon DuoHexaBody-CD37 treatment in the presence of Fc-crosslinker (Figure 4B, Supplementary Figure 4C). Similarly, ERK phosphorylation was upregulated in primary B cells upon DuoHexaBody-CD37 treatment in the presence of Fc crosslinker, in contrast to DLBCL cells (Figure 4B, Supplementary Figure 4C). Taken together, DuoHexaBody-CD37 treatment activates different signaling cascades in primary B cells versus DLBCL-derived tumor cells.

**Figure 4:**
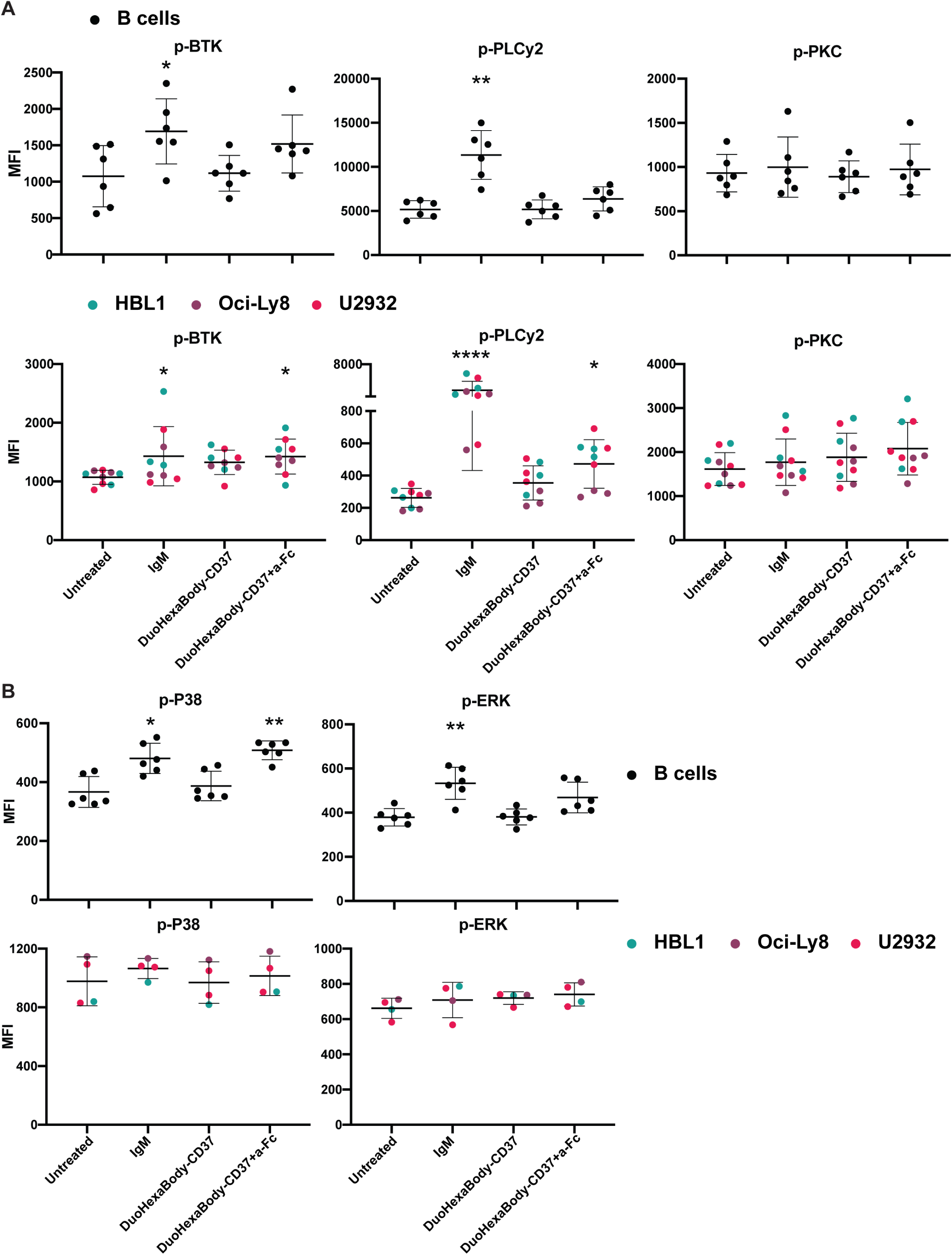
DuoHexaBody-CD37 treatment results in differential activation of BCR and RAS/MAPK downstream signaling proteins in primary B cells and DLBCL cell lines. (**A, B**) Phosphoflow analysis of **(A)** p-BTK(Y223), p-PLCy2(Y759), p-PKCα/βII(T638/641) and **(B)** p38(T180/Y182), p-ERK-1/2(T202/Y204) on primary B cells (*top row*) or DLBCL cells (*bottom row*) either untreated or treated with anti-BCR (F(ab’)2 anti-IgM), DuoHexaBody-CD37 and/or goat F(ab’)2 anti-human IgG (a-Fc). Dot plots depict quantification (mean +/- SD) of mean fluorescence intensity (MFI). Each dot represents an individual donor (primary B cells) or experimental replicate (DLBCL cell lines). Significance was calculated compared to untreated control using Kruskal-Wallis test followed by Dunn’ multiple comparison test correction (*p<0.05, **p<0.01, ****p<0.0001).

### CD37 N-terminus is involved in DuoHexaBody-CD37-mediated signaling

As tyrosine phosphorylation of the CD37 N-terminus was reported to be crucial in CD37 mediated signaling ^27^, we examined the involvement of CD37 N-terminus in DuoHexaBodyCD37-induced signaling. In these experiments, B-ALL cells (NALM6) with low endogenous CD37 expression were used to investigate various CD37 mutant constructs ^27^. NALM6 cells transfected with wild type CD37 (CD37-WT), or with CD37 carrying either the Tyr13 to phenylalanine mutation (CD37-Y13F) or a deletion of Tyr13 residue (CD37-ΔY13) in the cytosolic region of the CD37 molecule were generated. Cell surface expression of CD37-WT and both mutant variants was validated by flow cytometry (Figure 5A). DuoHexaBody-CD37 treatment in the presence of Fc-crosslinker resulted in significant upregulation of p-AKT(S473) and p-SHP1(Y564) in NALM6 cells expressing CD37-WT compared to treatment with only the crosslinker (Figure 5B, C) in line with findings observed in the DLBCL-derived cell lines (Figure 3C). Interestingly, deleting Tyr13 (CD37-ΔY13) completely impaired p-AKT(S473) and p-SHP1(Y564) upregulation, whereas mutating Tyr13 (CD37-Y13F) did not result in major changes upon DuoHexaBody-CD37 treatment in the presence of Fc-crosslinker (Figure 5B, C). In conclusion, these data validate active CD37 downstream signaling upon target engagement with DuoHexaBody-CD37 in the presence of Fc-crosslinker, and suggest the involvement of the CD37 N-terminus in mediating direct cytotoxic signaling. Since p-SHP1 upregulation was observed specifically in DuoHexaBody-CD37-treated cells upon Fc crosslinking, and SHP1 signaling was abolished when CD37-Y13 was mutated, we next investigated whether SHP1 knockout in Oci-Ly7 and HBL1 cells (Supplementary Figure 5A) affected the induction of direct cytotoxicity. No change in cytotoxicity was observed in either of the two cell lines following knockout of SHP1 (Supplementary Figure 5B), indicating that SHP1 signaling induced by DuoHexaBody-CD37 Fc-crosslinking does not directly contribute to the enhanced cytotoxicity.

**Figure 5:**
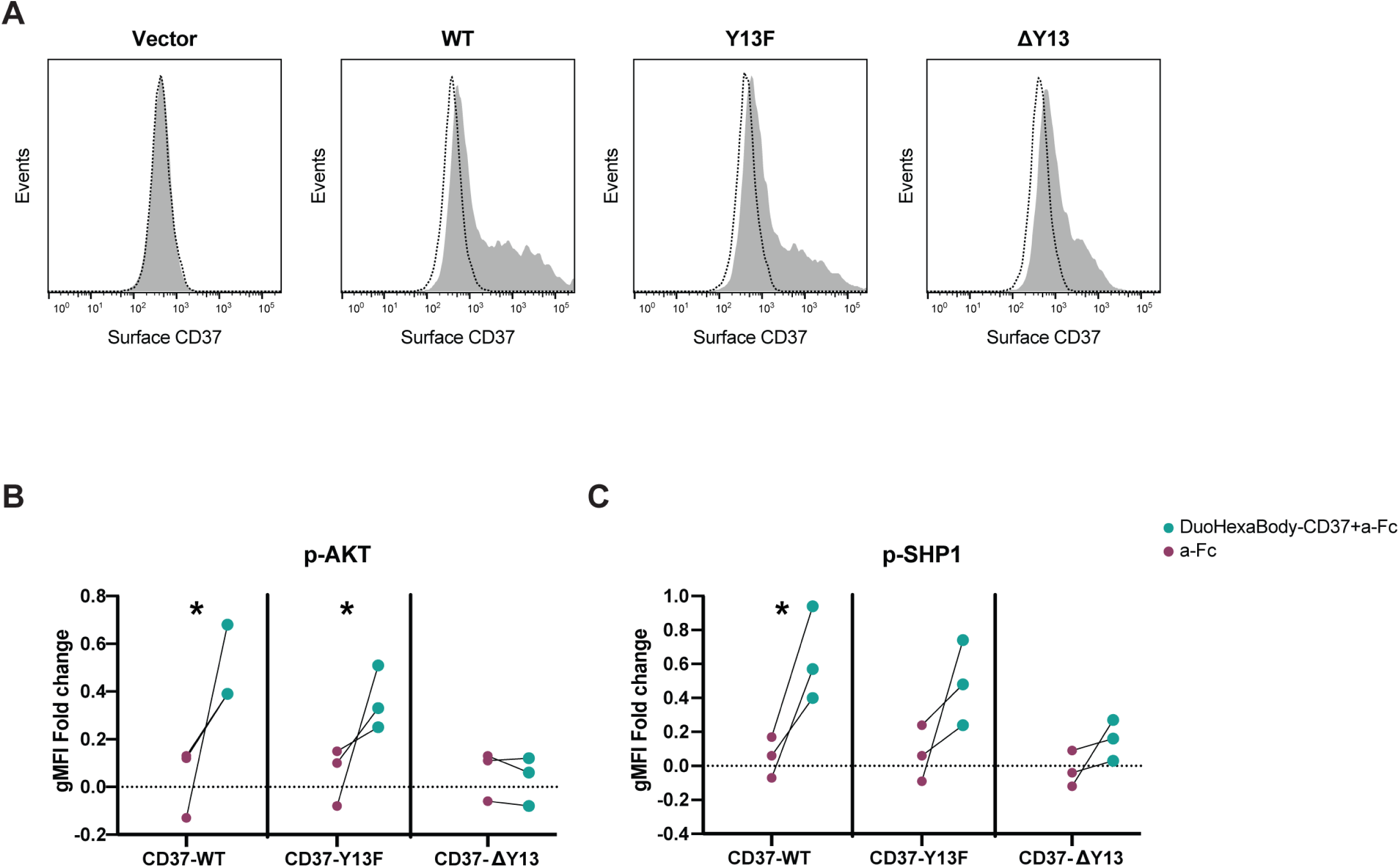
CD37 N-terminus is involved in DuoHexaBody-CD37-mediated cytotoxic signaling. (**A**) Histogram showing CD37 cell surface expression determined by flow cytometry analysis of NALM6 cell lines expressing wild type CD37 (CD37-WT), or mutations of Tyr13 to phenylalanine (CD37-Y13F), or deletion of Tyr13 (CD37-ΔY13) in the cytosolic regions of the CD37 molecule. (**B**) p-AKT(S473) and (**C**) p-SHP1(Y564) on NALM6 cell lines transfected with CD37-WT, CD37-Y13F or CD37-ΔY13 treated with DuoHexaBody-CD37 (DHB) and goat F(ab’)2 anti-human IgG (a-Fc) or crosslinker alone (a-Fc). Dot plots depict p-AKT and p-SHP1 levels in treated (DHB+a-Fc or a-Fc) vs untreated cells in CD37-GFP-positive (transfected) cells using phosphoflow analysis, corrected for background signal from GFP-negative cells. Significance was calculated using Student t-test (*p<0.05). Data shown from three independent experiments.

### DuoHexaBody-CD37 treatment abrogates cytokine mediated pro-survival signaling in DLBCL cells

As cells within the DLBCL tumor microenvironment can stimulate tumor growth by supplying cytokines ^38^ and CD37 has been reported to inhibit IL-6 signaling ^17^, we next investigated the impact of DuoHexaBody-CD37 treatment on cytokine-mediated oncogenic signaling. Different DLBCL cell lines were treated with DuoHexaBody-CD37 in absence or presence of Fc-crosslinker, followed by stimulation with either recombinant human IL-4 (rh-IL4), IL-6 (rhIL6) or IL-21 (rh-IL21) and analyzed for p-STAT activation by phosphoflow. These cytokines were selected because of their pro-tumorigenic role in DLBCL and other hematological cancers ^18, 38,39,40,41^. All tested DLBCL-derived tumor cell lines showed a significant increase in the level of p-STAT6 upon rh-IL-4 stimulation. Pre-treatment with DuoHexaBody-CD37 in the presence of a crosslinker showed a significant decrease in percentage p-STAT6-positive tumor cells upon stimulation with rh-IL-4 (Figure 6A) in contrast to controls (untreated or cells treated with DuoHexaBody-CD37 or Fc-crosslinker alone). Next, IL-6-induced p-STAT3 signaling was investigated in HBL-1 cells, as the other DLBCL-derived tumor cell lines did not respond to IL-6 (data not shown). A two-fold increase in percentage p-STAT3-positive HBL-1 cells was observed upon rh-IL6 stimulation. Interestingly, treatment with DuoHexaBody-CD37 in the presence of a crosslinker also decreased the percentage of p-STAT3-positive HBL-1 cells compared to controls (untreated or cells treated with DuoHexaBody-CD37 or crosslinker alone) although this was not significant (Figure 6B). Finally, IL-21-induced p-STAT3 signaling was investigated in all four cell lines. Both ABC-DLBCL derived tumor cells (HBL-1 and U2932) showed a > four-fold increase in the percentage of p-STAT3-positive cells, while the effect was more modest in the GCB-DLBCL-derived tumor cells (Oci-Ly8) showing a two-fold increase in p-STAT3 signal. Oci-Ly7 cells did not respond to IL-21 (data not shown). DuoHexaBody-CD37 treatment in the presence of Fc-crosslinker showed in all three IL-21 responsive cell lines a significant decrease in the percentage of p-STAT3-positive cells upon rh-IL21 stimulation (Figure 6C) in line with IL-4 and IL-6 stimulation. Collectively, these experiments show that DuoHexaBody-CD37 treatment in the presence of Fc-crosslinker inhibits cytokine-mediated pro-survival signaling in the DLBCL tumor microenvironment.

**Figure 6:**
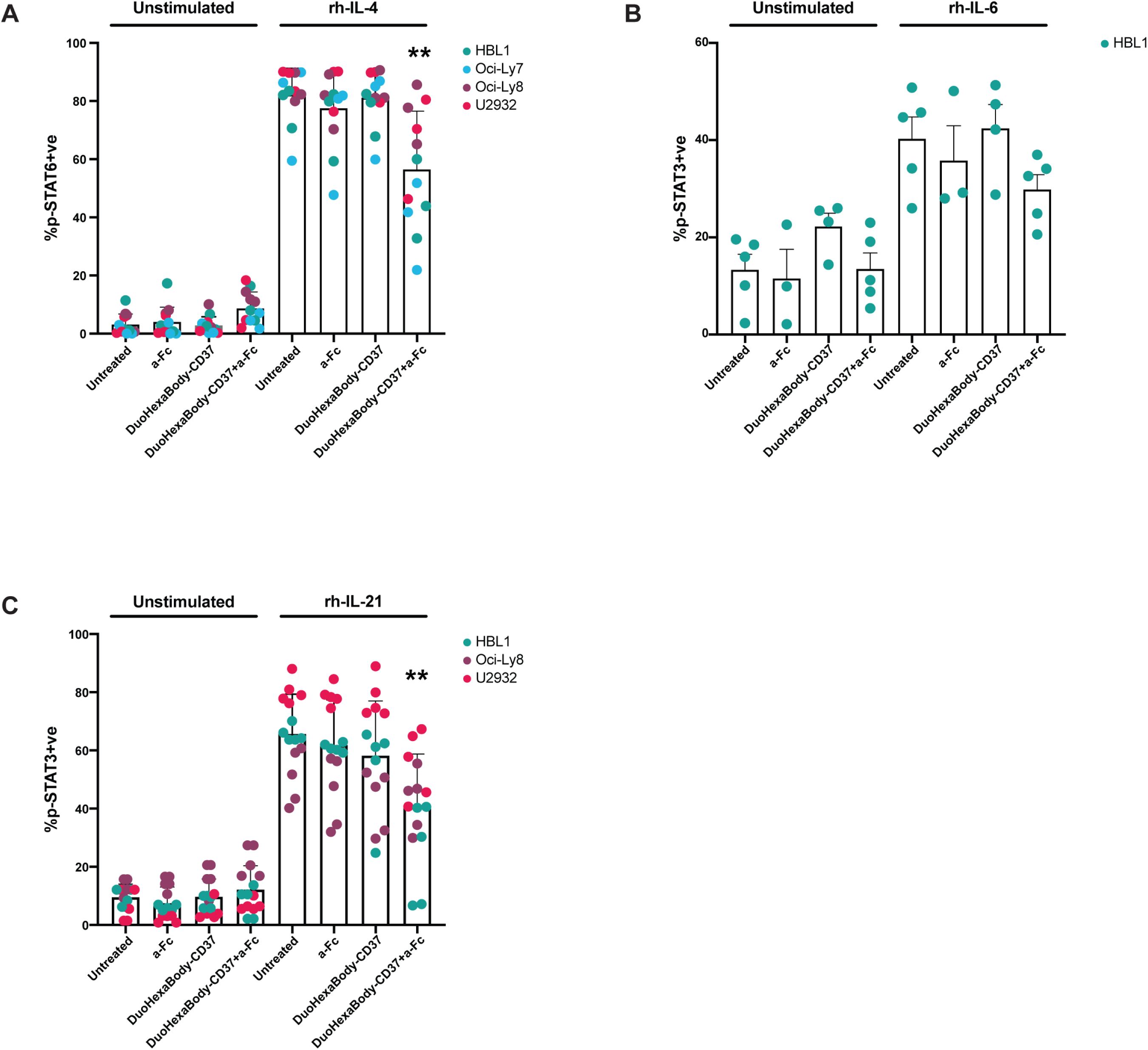
DuoHexaBody-CD37 treatment decreases cytokine-mediated pro-survival signaling in DLBCL cell lines. (**A**) Percentages of p-STAT6-positive cells (mean and SD) upon recombinant human IL-4 (rh-IL4) stimulation in DLBCL cells pre-treated with DuoHexaBody-CD37 in presence or absence of goat F(ab’)2 anti-human IgG (a-Fc). Data is shown from at least three independent experiments. Percentages of p-STAT3-positive cells (mean +/- SD) upon (**B**) recombinant human IL-6 (rh-IL-6) stimulation in HBL-1 cells (the only IL-6 responsive cell line in this study) or (**C**) recombinant human IL-21 (rh-IL-21) stimulation in DLBCL cells pre-treated with DuoHexaBody-CD37 in presence or absence of a-Fc. Data is shown from at least three independent experiments. (**A-C**) Significance was calculated compared to untreated control using Kruskal-Wallis test followed by Dunn’ multiple comparison test correction (**p<0.01).

## Discussion

A large fraction (∼40%) of patients with DLBCL do not respond to or relapse after first-line treatment with standard immunochemotherapy (R-CHOP), emphasizing the unmet medical need to develop new therapies for DLBCL. Tetraspanins have gained strong interest as therapeutic targets in different cancer types due to their capacity to regulate cell proliferation, adhesion and migration. Importantly, many tumor cells have altered cell surface expression of tetraspanins making them attractive targets for the treatment of cancer ^16,42^. For example, CD151 is upregulated in carcinomas and CD151 stimulates both primary tumor growth and the metastatic cascade through its interaction with β1 integrins ^42,43^. In B cell lymphoma, CD81 has been reported to be highly expressed by malignant cells from patients with B-NHL, and CD81-specific antibodies have been reported to be cytotoxic for such cells while sparing normal B cells ^44^. CD37 represents an alternative therapeutic target for B-NHL because of its specific membrane expression on mature B cells, and its prognostic value in DLBCL and follicular lymphoma ^4,45^. DuoHexaBody-CD37 has been previously reported to mediate cytotoxicity of DLBCL cells via different mechanisms of action: complement (CDC) and effector cell-mediated killing (ADCC/ADCP) ^25,26^. Here, we report an additional mechanism of action of DuoHexaBody-CD37: CD37-mediated direct cytotoxicity which is independent of complement. Direct antibody-mediated killing may be particularly important in the DLBCL tumor microenvironment when complement is not available or inhibited. Complement has a dual role in tumors including hematological malignancies. The protective role of complement in antibody-mediated CDC has been well-established. However, more recent studies have provided evidence that tumors can hijack the complement components by cell surface expression of C1q, C3a and C5a receptors that promotes tumor growth and progression ^46,47^. In addition, lymphoma cells can upregulate complement regulatory proteins (CRPs: CD46, CD55 and CD59) to escape from antibody mediated CDC ^48^. Although no direct correlation was found between DuoHexaBody-CD37-induced CDC and expression of CRPs in vitro ^26^, expression of CD59 has been shown to predict clinical outcome in DLBCL patients treated with R-CHOP ^49^. In addition, the presence of regulatory myeloid cells (tumor-associated neutrophils, myeloid-derived suppressor cells) in the tumor microenvironment has been associated with immune suppression and poor prognosis in lymphoma (reviewed in ^38,50^). These tumor-infiltrating myeloid cells stimulate tumor growth and may impair ADCC/ADCP activity of therapeutic antibodies ^51^. Lastly, DLBCL cells have been shown to upregulate CD47 to protect themselves from ADCC and ADCP ^52^. Taken together, inhibition of complement activation by tumor cells or ineffective ADCC/ADCP may limit efficacy of antibody-based immunotherapies for hematological malignancies.

Since decreased expression of CD20 is related to inferior clinical outcome of DLBCL patients after treatment with R-CHOP, DuoHexaBody-CD37 treatment may be more potent compared to CD20-targeting therapies in DLBCL. Supporting this, we observed no/minimal internalization of CD37 by DuoHexaBody-CD37 in contrast to rituximab-induced CD20 internalization. We observed that DLBCL cells were more sensitive to DuoHexaBody-CD37mediated cytotoxicity than Burkitt cells, a difference that cannot be explained by variations in CD37 surface expression or clustering. It is possible that Burkitt cells rely on other oncogenic signaling pathways (i.e. MYC-driven) compared to DLBCL cells.

The activation of multiple intracellular signaling pathways, including PI3K-AKT and RAS-MAPK that play a central role in cancer, was observed upon DuoHexaBody-CD37 treatment of DLBCL cells. This was dependent on Fc-crosslinking of DuoHexaBody-CD37, which is in line with former studies using CD37-specific small modular immunopharmaceuticals (CD37-SMIP) ^53,54^. In vivo crosslinking could be facilitated by effector cells expressing Fc receptors, as evidenced by the observed direct cytotoxicity of DuoHexaBody-CD37 in tumor cells in the presence of PBMCs. This did not require FcγR mediated signaling or effector cell function (degranulation, trogocytosis) as fixed PBMCs could be used to facilitate the crosslinking. CD37 can activate and inhibit AKT kinase through its two intracellular tails that contain ITIM-like and ITAM-like motifs ^27^. Different mechanisms of direct tumor cell death by antibodies have been reported, including apoptosis, necrosis, pyroptosis and necroptosis. We identified apoptosis as the mechanism underlying DuoHexaBody-CD37–induced killing of DLBCL cells, and future studies, and future studies are needed to determine whether any of the other mechanisms are involved as well. mechanisms underlie DuoHexaBody-CD37-induced killing of DLBCL cells. Interestingly, increased p-AKT was observed in primary B cells upon DuoHexaBody-CD37 treatment in contrast to malignant B cells, indicating that primary B cells may be less sensitive to direct killing than lymphoma cells. Our results show that DuoHexaBody-CD37 directly affects the membrane-proximal signaling protein AKT which may provide new opportunities for combination therapies in DLBCL ^55^. For example, studies combining CD37-targeting antibodies with PI3K inhibitors ^56,57^ or cell cycle kinase inhibitors ^58^ demonstrated enhanced tumor cell death. We also observed p-BTK and p-PLCy upregulation in ABC-DLBCL upon DuoHexaBody-CD37 treatment, which could provide a rational to combine DuoHexaBodyCD37 with ibrutinib in ABC-DLBCL where BTK inhibition is already more effective compared to GBC-DLBCL ^59,60^. DuoHexaBody-CD37 treatment also led to an increase in SHP1 mediated signaling, however we could not confirm a direct role of SHP1 signaling in DuoHexaBody-CD37-mediated cytotoxicity. DLBCL cells may undergo signal rewiring upon SHP1 knockdown by altered levels of p-AKT, p-STAT3, and p-STAT6, or SHP2 may compensate for the loss of SHP1. It is currently unclear what the biological implications are of the increased SHP1 signaling observed upon treatment with DuoHexaBody-CD37 in DLBCL cells.

Moreover, DuoHexaBody-CD37 was found to be particularly effective in downregulating p-STAT3 and p-STAT6 in presence of IL-21 and IL-4, respectively. Within the DLBCL tumor microenvironment, these cytokines actively contribute to cancer pathogenesis in ABC-DLBCL ^18,38,39,40,41^. p-STAT3 downregulation upon DuoHexaBody-CD37 treatment in presence of IL-6 requires further investigation in additional IL-6-responsive cell lines, as HBL1 was the only IL-6-responsive lymphoma cell line tested in this study. IL-6 levels in serum are associated with adverse clinical outcome in DLBCL, and we previously reported that CD37 inhibits IL-6 receptor signaling through SOCS3 ^17^. In line with this, patients with CD37-negative DLBCL present with higher IL-6 levels in serum and tumors which is associated with inferior clinical outcome ^17^. CD37 is an independent prognostic factor in DLBCL, irrespective of DLBCL subset ^4^, which has been confirmed in patients with follicular lymphoma ^45^. We anticipate that targeting CD37, especially in the presence of classical chemotherapy and/or targeted drugs will further enhance the efficacy of immunotherapy of DLBCL. In conclusion, this study shows DuoHexaBody-CD37 induces cytotoxic signaling in DLBCL cells, and reduces cytokine-mediated pro-survival signaling in the DLBCL tumor microenvironment.

## Material and Methods

### Therapeutic antibodies

DuoHexaBody-CD37 (GEN3009) and the negative control anti-HIV-1 gp120 antibody IgG1-b12 (mentioned in manuscript as IgG1-ctrl) were generated by Genmab (Utrecht, The Netherlands) as previously described ^61^. Rituximab anti-CD20 (Mab Thera®) was obtained from Radboudumc (Nijmegen, the Netherlands).

### Cell culture

Lymphoma cell lines (GCB-DLBCL: Oci-Ly8, Oci-Ly7; ABC-DLBC: HBL-1, U2932; B-ALL: NALM6, and Burkitt: BJAB, DAUDI) were obtained from DSMZ, ATCC, Blanca Scheijen (Dept. Pathology, Radboudumc) and Marcel Spaargaren (Dept. of Pathology, AmsterdamUMC) and cell lines were authenticated using STR-analysis. Cells were cultured in RPMI 1640 (Gibco), 1% antibiotic-antimycotic (Gibco) and 10% FBS (Hyclone) (RPMI/10%FCS) and incubated at 37°C with 5% CO_2_. For optimum growth, the cell lines were propagated in dilution of 0.5 × 10^6^ cells/ml and the culture media were refreshed twice a week. All experiments were performed one day after refreshing medium and expanding cell culture.

### MTS assay

Cell viability was determined using CellTiter 96® AQueous One Solution Reagent from Promega as per manufacturer instructions. Briefly, 2x10^5^ cells in RPMI/10%FCS were plated in 96-well plates and treated with 5µg/ml DuoHexaBody-CD37 for 30min at 4°C. Cells were washed and treated with 5 µg/ml of goat F(ab’)2 anti-human IgG (a-Fc, Southern Biotech) and incubated for 48h at 37°C with 5% CO_2_. As control, cells treated with DuoHexaBody-CD37 or a-Fc alone were included. Subsequently, cells were stained with 1:5 dilution of CellTiter 96® AQueous One Solution Cell Proliferation Assay (MTS, Promega) in media (RPMI/10%FCS) and incubated at 37°C with 5% CO_2_ for 2-3hrs. Absorbance was recorded at 490nm in a 96-well plate reader (Bio-Rad iMark Microplate Absorbance Reader).

### Cytotoxicity assays

Peripheral blood from healthy volunteers was obtained via Sanquin blood bank (Nijmegen, NL) upon informed consent and anonymized for further use, following the guidelines of the Institutional Review Board, and in accordance with the declaration of Helsinki. Peripheral blood mononuclear cells (PBMCs) were isolated using Ficoll Hypaque (GE Healthcare, Little Chalfont, UK) according to the manufacturer’s instructions. For studying different subsets of FcγR-expressing cell types, B cell, T cells, monocytes and NK cells were isolated using Miltenyi Biotec cell separation reagents (130-101-638, 130-096-495, 130-117-337 and 130-092-657, respectively). For fixation, PBMCs and the purified immune cell subsets were washed with PBS and resuspended directly in 1%PFA (filter sterilized) at 7.5x10^6^ cells/ml for 20min at RT (room temperature). Following, cells were washed twice in PBS and resuspended in media (RPMI/10%FCS). In parallel, 10x10^6^ target cells (HBL-1 or U2932) were stained with CellTrace Violet Cell Proliferation Kit (Invitrogen) at final concentration of 167nM for 20min at 37°C with 5% CO_2_. Subsequently, cells were washed twice and resuspended in media (RPMI/10%FCS). Next, CTV-stained target cells were treated with 5µg/ml DuoHexaBody-CD37 for 30min at 4°C and washed with media. In addition, target cells were co-cultured with fixed PBMCs at 1:20 (target to crosslinker) ratio for 4, 24 or 48h (or for 3 hours with the purified immune cell subsets), at 37°C in the presence of 5% CO_2_. Subsequently, cells were directly stained with 5µg/ml propidium iodide (Miltenyi Biotec) for 5min at RT on plate shaker and analyzed using FACS Lyric (BD Biosciences).

### Target binding analysis

Briefly, 100,000 lymphoma cells (HBL-1, Oci-Ly7, U2932, BJAB or DAUDI) were treated with 5mg/ml of anti-CD20 Rituximab (Mab Thera®) or DuoHexaBody-CD37 in RPMI/0.2%BSA for 15min at RT in a 96-wells plate. Plates were then directly transferred to incubator (37°C/5% CO_2_) for 0, 2 or 6h. Cells were harvested at respective time points by washing with PBA buffer (PBS, 1% BSA, azide 0.02%) followed by staining with mouse anti-human IgG1 Fc secondary antibody conjugated to Alexa Fluor 488 (ThermoFisher) for 30min at 4°C. Next, cells were washed and stained with Live-dead marker (eFluor 506, eBioscience) and measured using FACS Lyric (BD Biosciences).

### Generation of CD37 mutant cell lines

NALM6 cells were used because of their endogenous low CD37 expression. Cells were grown in RPMI/10% FBS supplemented with glutamine 2mM. Cells were transfected with either CD37-sGFP2, CD37 Y13F-sGFP2 or CD37-delta Y13-sGFP2 constructs using the Amaxa 4D nucleofector. Briefly, for each transfection 3x10^6^ cells were collected and washed with PBS. Cells were resuspended in 100 µl SF buffer (cat. no. V4XC-2024) containing 3 µg of plasmid and pulsed with program CV-104. After transfection, cells were transferred to 6-well plates and incubated overnight in 3 ml of RPMI with 10% FBS without antibiotic-antimycotic and phenol red. The CD37 constructs were kindly provided by Prof. Dr. N. Muthusamy (Ohio University, USA) ^27^, and subcloned in pSGFP-N1 vector using standard techniques. Sequences were verified via sequencing.

### Phosphoflow assays

Lymphoma cells (2x10^5^ cells/well) or PBMCs (5x10^5^ cells/well) in RPMI/10%FCS were treated with 5µg/ml DuoHexaBody-CD37 for 30min at 4°C. Next, cells were washed and treated with 5µg/ml of goat F(ab’)2 anti-human IgG (a-Fc) and incubated for 20min at 37°C with 5% CO_2_. As treatment control, cells were treated with DuoHexaBody-CD37 or a-Fc alone. As positive control of B-cell receptor stimulation, cells were treated with 20µg/ml of goat F(ab’)2 antihuman IgM. Next, cells were spun down and stained at 1:10 dilution with Fixable Viability stain 510 (BD Biosciences) in RPMI/10%FCS for 10min at 37°C. Thereafter, cells were directly fixed for 10min at 37°C with 100 μl of the eBioscience FoxP3/Transcription Factor Fixation kit (Invitrogen) as indicated by the manufacturer. After fixation, samples were washed twice with eBioscience Permeabilization Wash Buffer (10X diluted in Milli-Q) (Invitrogen). To identify the B cells within PBMC cultures, cells were stained with antibodies against CD19, CD3 and IgD diluted in eBioscience wash buffer. To prevent non-specific antibody binding, FcR-blocking purified rat anti-mouse CD16/CD32 (Mouse BD Fc BlockTM) was included in the staining. Stained cells were incubated for 30min at 4°C followed by washing in eBioscience wash buffer and staining with antibodies against phosphorylated proteins; p-SHP1(Y564), p-AKT(S473), p38(T180/Y182) (all Cell signaling technology), p-SYK(Y348), p-BTK(Y223), pPLCy2(Y759), p-PKCα/βII(T638/641), p-ERK-1/2(T202/Y204) (all BD Biosciences); in wash buffer for 30min at RT. Unlabeled phospho-antibodies (p-SHP1(Y564), p-AKT(S473), p38(T180/Y182)) were stained with a secondary PE-labeled F(ab’)2 donkey anti-rabbit IgG (H+L) antibody (Jackson Immuno Research) for 15min at RT. Lastly, the cells were washed and resuspended in wash buffer before measuring on FACS Lyric.

### Cytokine stimulation experiments

To measure p-STAT3 and p-STAT6 responses, cells treated with DuoHexaBody-CD37 and/or goat F(ab’)2 anti-human IgG (a-Fc) for 24h as described above. After washing, cells were stimulated with recombinant IL-6 (100ng/ml) or IL-4 (20ng/ml) (both from Miltenyi) or IL-21 (Peprotech) (50ng/ml) for 15min at 37°C. Cells were harvested and stained with Fixable Viability stain 510 (BD Biosciences). Subsequently, cells were fixed with BD Cytofix Fixation Buffer (BD Biosciences) for 10min at 37°C and washed twice with MACS buffer (PBS/0.5% BSA/2 mM EDTA). Next, cells were permeabilized using 150 μl Perm Buffer III (BD Biosciences) at 4°C for 30min and washed twice with MACS buffer, followed by staining with PE-labeled p-STAT3 (Y705) or p-STAT6 (Y641) (BD Biosciences) antibody diluted (1:8) in MACS buffer for 30min at RT. Finally, the cells were washed and resuspended in 50 μl MACS buffer before measuring on FACS Lyric.

## Supplementary methods

Microscopy analysis of CD37 clustering, RPPA assay, DuoHexaBody-CD37 surface accessibility assays, CRISPR-cas9 knockout of SHP1 in DLBCL cell lines and quantification of apoptosis.

## Acknowledgements

We thank the Radboudumc Microscopy Imaging Center for use of their microscopy facilities, as well as for their support and assistance. We thank Simone Oostindie and Inge Verbrugge for valuable scientific input, and Peter Friedl for facilitating the RPPA studies. The CD37 constructs were kindly provided by Natarajan Muthusamy (Ohio University, USA) ^27^.

## Author contributions

ABvS, SPS, ECWB, MGR, KCMS and MBO provided scientific input and conceptualized the study. SPS developed the methodology and performed the data analysis for the cytotoxicity assays, PBMC assays, phosphoflow experiments, CD37 mutant studies, cytokine studies and RPPA analysis. KM performed FcγR cytotoxicity studies, CD37 expression studies and apoptosis assays. MDvdB performed the surface accessibility assays and CD37 mutant studies. SvD developed the methodology for the CD37 transfections. MTB developed methodology of CD37 microscopy studies and performed data analysis on CD37 clustering. WC generated and validated the SHP-1 KO cells and performed cytotoxicity assays. SPS and ABvS wrote the manuscript. ABvS supervised the study. All authors read, revised, and agreed on the final version of the manuscript.

---

**Supplementary Figure 1:**
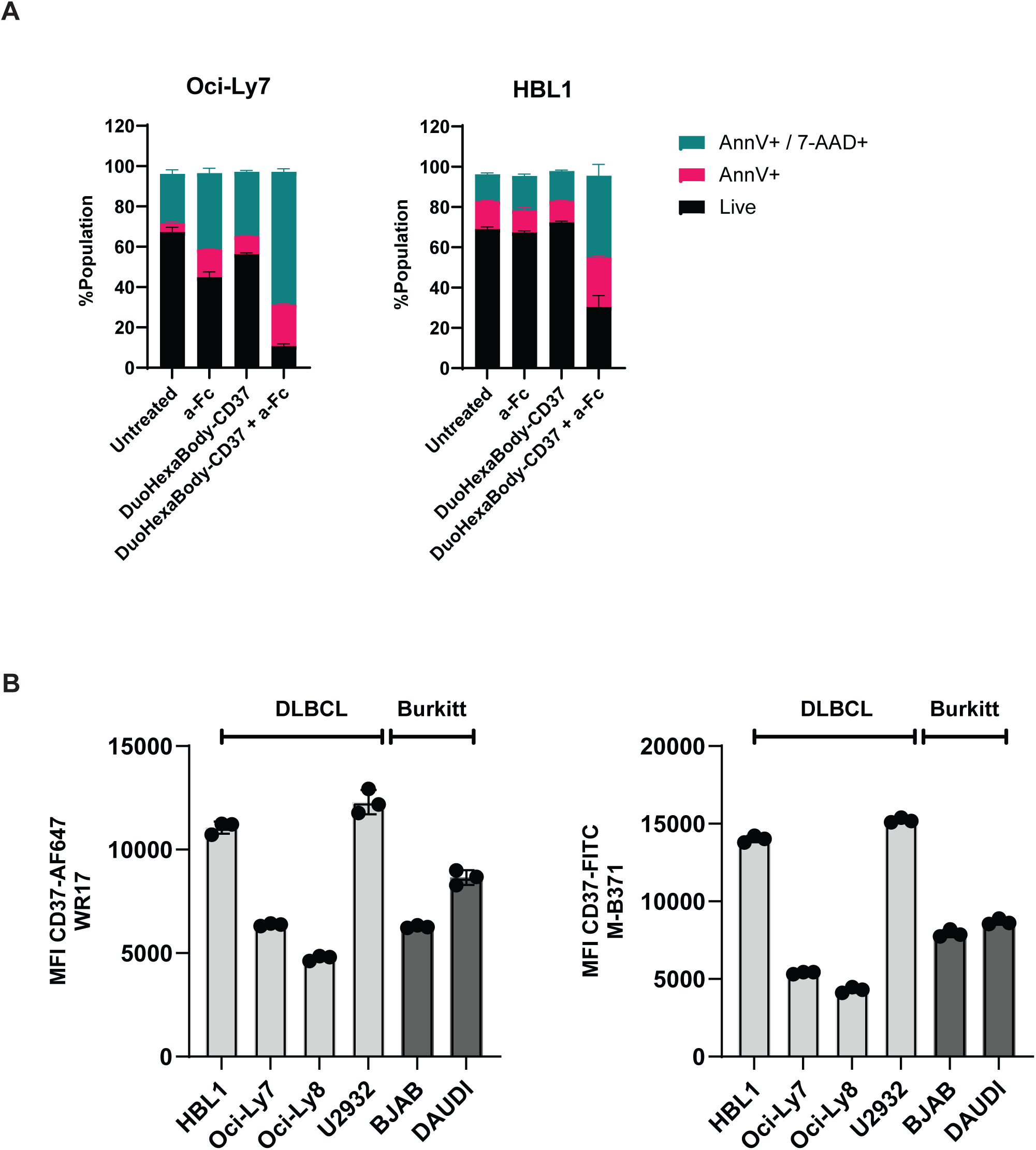
Fc-crosslinking-mediated apoptotic cell death induced by DuoHexaBody-CD37 is not correlated with CD37 membrane expression. **(A)** Flow cytometric comparison of Annexin V stained, Annexin V + 7AAD stained and unstained cells in presence or absence of DuoHexaBody-CD37, with and without Fc-crosslinking on Oci-Ly8, Oci-Ly7 and HBL 1 cell lines. Data (mean +/- SD) show unstained, Annexin V-positive and Annexin V + 7AAD double positive populations from three independent experiments. **(B)** Flow cytometric quantification of CD37 expressed on membranes of DLBCL and Burkitt cell lines using two different CD37 antibodies (clones WR-17 and M-B371). Data (mean +/- SD) represent three independent experiments. MFI=mean fluorescence intensity.

**Supplementary Figure 2:**
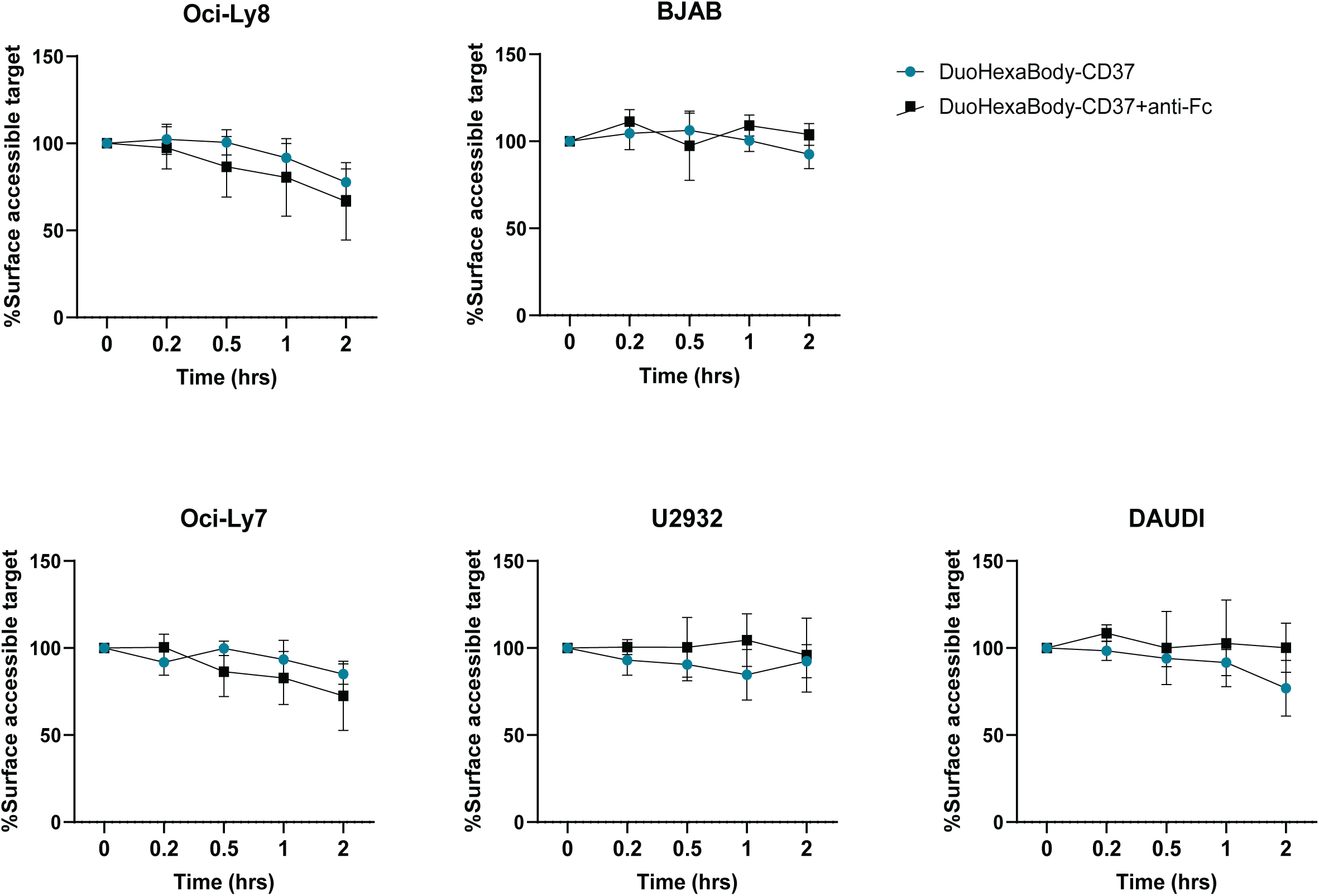
Fc-crosslinking does not lead to increased CD37 internalization from the cell surface of DLBCL cell lines upon DuoHexaBody-CD37 treatment. Percentage decrease in cell surface binding by DuoHexaBody-CD37 with or without Fc-crosslinker in Oci-Ly8, BJAB, Oci-Ly7, U2932 and DAUDI cell lines as measured by flow cytometry. Data (mean +/- SD) is derived from four independent experiments.

**Supplementary Figure 3:**
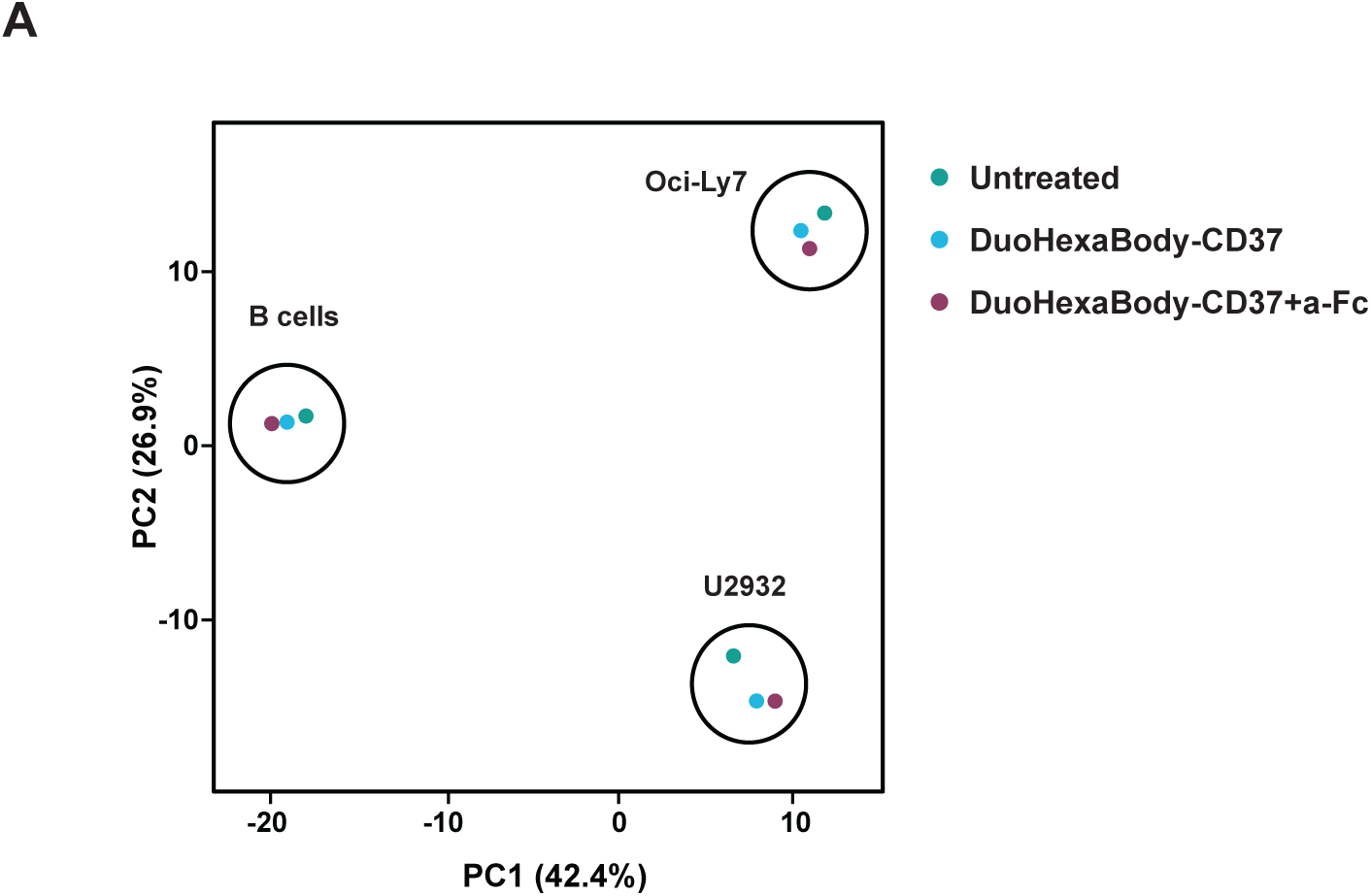
Principal component analysis of (phospho)protein expression profiles after DuoHexaBody-CD37 treatment. Principal component analysis (PCA) of 484 proteins (102 phospho targets and 382 total proteins) in primary B cells (CD19+ purified) and two DLBCL cell lines (Oci-Ly7 and U2932) either untreated or treated with DuoHexaBody-CD37 in presence or absence of a-Fc using reverse phase protein array (RPPA). The first two components of the cluster analysis are shown.

**Supplementary Figure 4:**
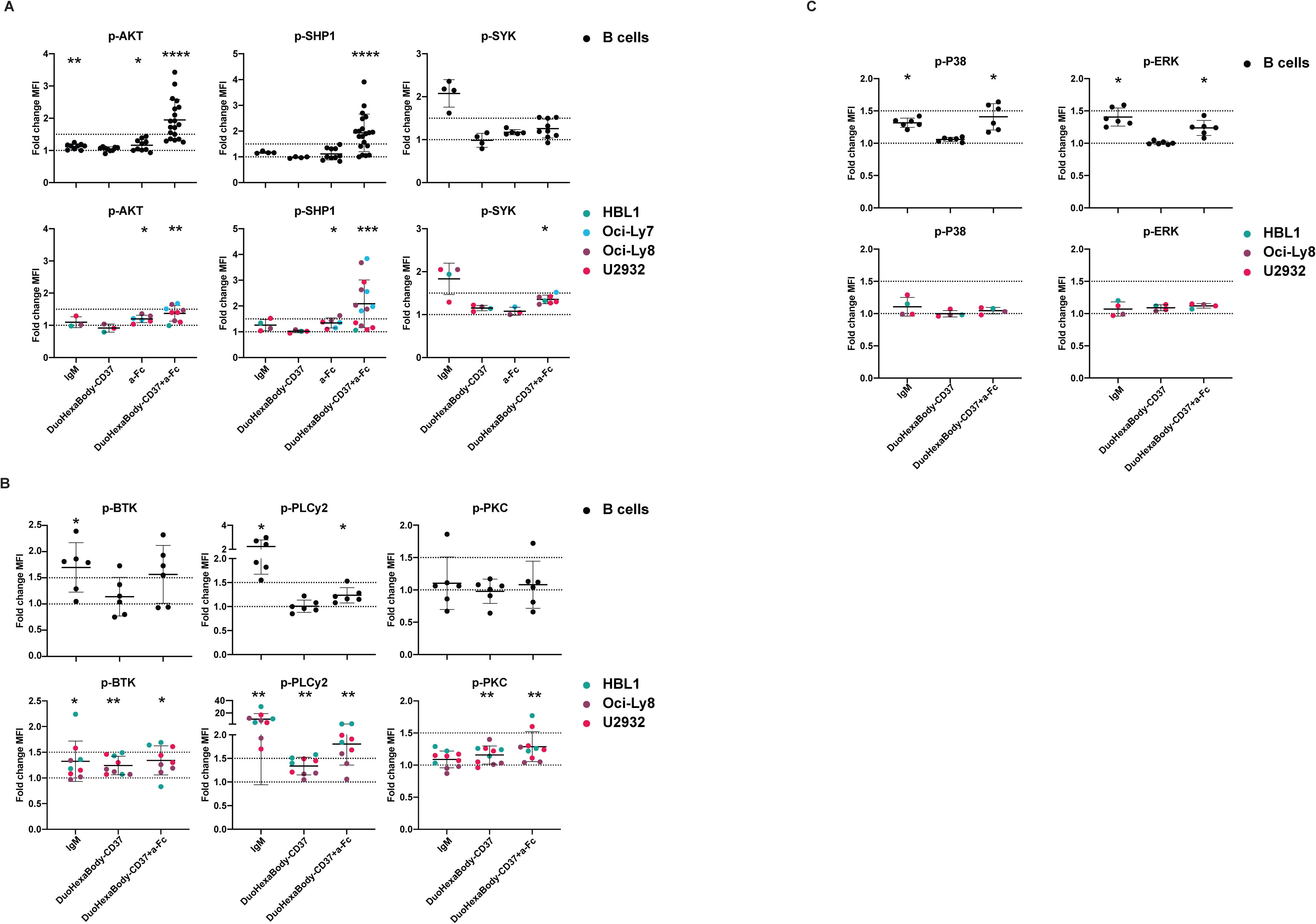
DuoHexaBody-CD37 treatment results in differential activation of BCR and RAS/MAPK downstream signaling proteins in B cells and DLBCL. **(A-C)** Phosphoflow analysis of (A) p-SYK(Y348), p-SHP1(Y564), p-AKT(S473); **(B)** p-BTK(Y223), p-PLCy2(Y759), p-PKCα/βII(T638/641) and **(C)** p38(T180/Y182), p-ERK-1/2(T202/Y204) on primary B cells (*top row*) or DLBCL cells (*bottom row*) either anti-IgM treated or treated with DuoHexaBody-CD37 and/or goat F(ab’)2 anti-human IgG (a-Fc). Dot plots depict fold change (mean +/- SD) of mean fluorescence intensity (MFI) of indicated treatment to untreated cells. Each dot represents individual donor (B cells) or experimental replicate (DLBCL). Significance of fold change to untreated control was calculated using Wilcoxon Signed Rank test (*p<0.05, **p<0.01, ****p<0.0001).

**Supplementary Figure 5:**
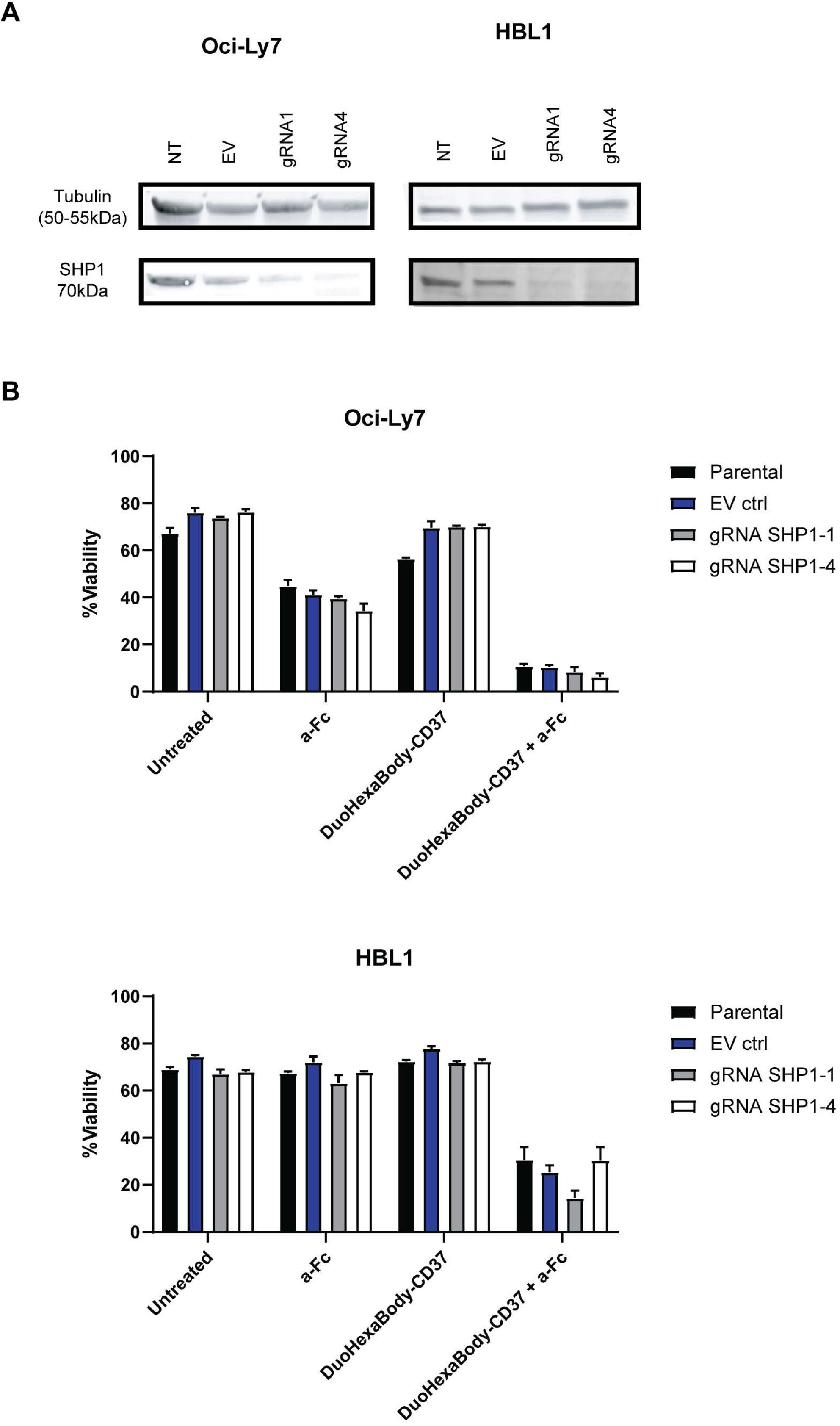
Knockout of SHP1 does not influence DuoHexaBody-CD37 mediated cytotoxicity. **(A)** Western blot showing SHP-1 levels in Oci-Ly7 and HBL1 cells for parental (NT), empty vector (EV) and SHP-1 gRNA1/gRNA4 transduced cells. Tubulin was used as loading control. (B) Percentage of dead (Annexin V + 7AAD double positive) populations in parental, transduction control (gRNA EV), and SHP1 knockout (gRNA SHP1-1 and gRNA SHP1-4) in Oci-Ly7 and HBL1 cells measured by flow cytometry. Data (mean +/- SD) shown from three independent experiments.

**Supplementary Table 1.**
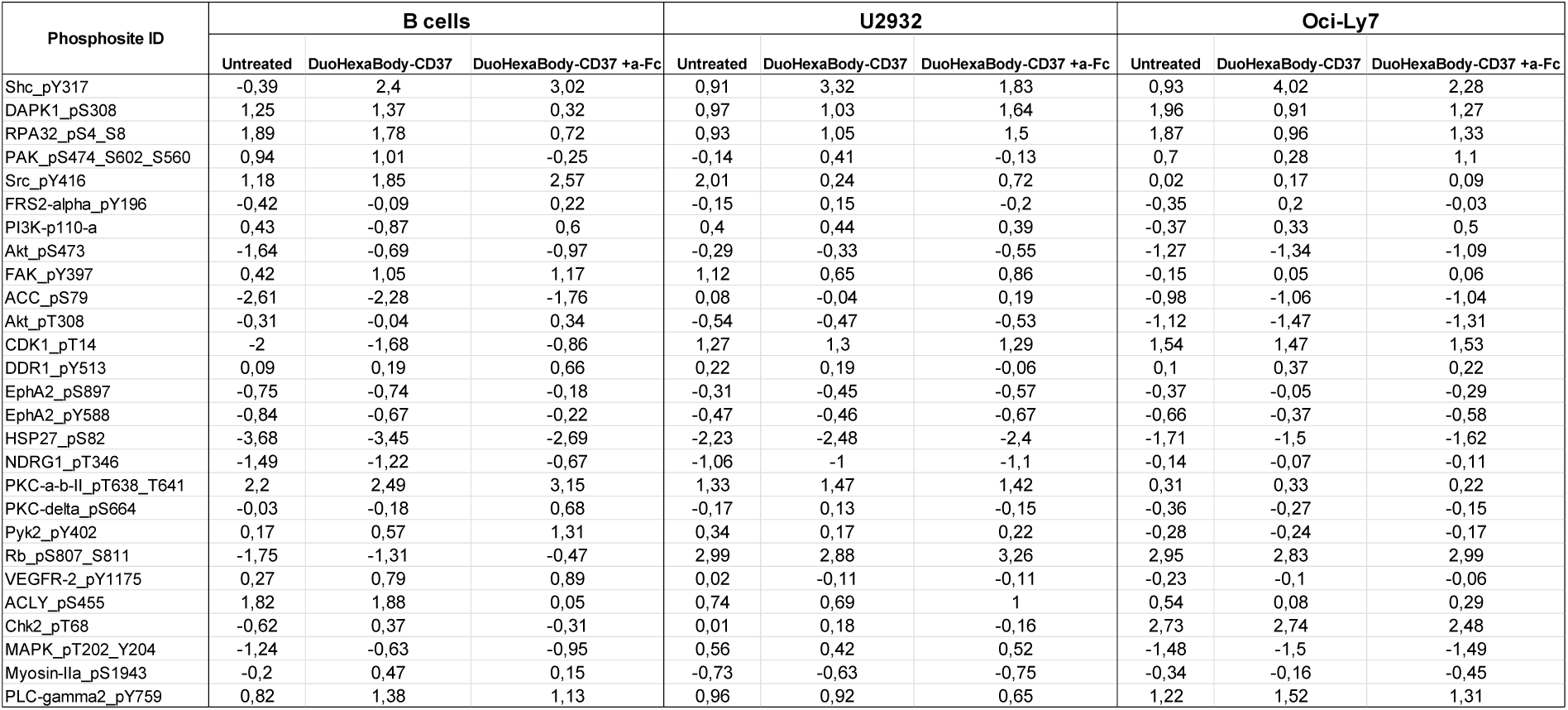

## References

1. Morin RD, Arthur SE, Hodson DJ. Molecular profiling in diffuse large B-cell lymphoma: why so many types of subtypes? Br J Haematol. 2022;196(4):814–29.

2. Abou Dalle I, Dulery R, Moukalled N, Ricard L, Stocker N, El-Cheikh J, et al. Bi- and Trispecific antibodies in non-Hodgkin lymphoma: current data and perspectives. Blood Cancer J. 2024;14(1).

3. Johnson NA, Boyle M, Bashashati A, Leach S, Brooks-Wilson A, Sehn LH, et al. Diffuse large B-cell lymphoma: Reduced CD20 expression is associated with an inferior survival. Blood. 2009;113(16):3773–80.

4. Xu-Monette ZY, Li L, Byrd JC, Jabbar KJ, Manyam GC, De Winde CM, et al. Assessment of CD37 B-cell antigen and cell of origin significantly improves risk prediction in diffuse large B-cell lymphoma. Blood. 2016;128(26):3083–100.

5. Guan XW, Wang HQ, Ban WW, Chang Z, Chen HZ, Jia L, et al. Novel HDAC inhibitor Chidamide synergizes with Rituximab to inhibit diffuse large B-cell lymphoma tumour growth by upregulating CD20. Cell Death Dis. 2020;11(1).

6. Roschewski M, Staudt LM, Wilson WH. Diffuse large B-cell lymphoma—treatment approaches in the molecular era. Nat Rev Clin Oncol. 2013;11(1):12–23.

7. Varma G, Goldstein J, Advani RH. Novel agents in relapsed/refractory diffuse large B cell lymphoma. Hematol Oncol. 2023;41(S1):92–106.

8. Schwartz-Albiez R, Dörken B, Hofmann W, Moldenhauer G. The B cell-associated CD37 antigen (gp40-52). Structure and subcellular expression of an extensively glycosylated glycoprotein. J Immunol 1988;140(3):905–14.

9. Van Spriel AB, De Keijzer S, Van Der Schaaf A, Gartlan KH, Sofi M, Light A, et al. The tetraspanin CD37 orchestrates the α4β1 integrin-Akt signaling axis and supports long-lived plasma cell survival. Sci Signal. 2012;5(250):ra82.

10. de Winde CM, Zuidscherwoude M, Vasaturo A, van der Schaaf A, Figdor CG, van Spriel AB. Multispectral imaging reveals the tissue distribution of tetraspanins in human lymphoid organs. Histochem Cell Biol. 2015; 144(2):133–46.

11. Berditchevski F, Rubinstein E. Tetraspanins. 2013. 1–418 p. http://link.springer.com/10.1007/978-94-007-6070-7.

12. Hemler ME. Targeting of tetraspanin proteins — potential benefits and strategies. Nat Rev Drug Discov. 2008;7(9):747–58.

13. Querol Cano L, Dunlock VME, Schwerdtfeger F, van Spriel AB. Membrane organization by tetraspanins and galectins shapes lymphocyte function. Nat Rev Immunol. 2024;24(3):193–212.

14. Barrena S, Almeida J, Yunta M, Lopez A, Fernandez-Mosteirin N, Giralt M, et al. Aberrant expression of tetraspanin molecules in B-cell chronic lymphoproliferative disorders and its correlation with normal B-cell maturation. Leukemia. 2005;19(8):1376–83.

15. Bobrowicz M, Kubacz M, Slusarczyk A, Winiarska M. Cd37 in b cell derived tumors— more than just a docking point for monoclonal antibodies. Int J Mol Sci. 2020;21(24):1.

16. Beckwith KA, Byrd JC, Muthusamy N. Tetraspanins as therapeutic targets in hematological malignancy: A concise review. Front Physiol. 2015 Mar 23;6.

17. De Winde CM, Veenbergen S, Young KH, Xu-Monette ZY, Wang XX, Xia Y, et al. Tetraspanin CD37 protects against the development of B cell lymphoma. J Clin Invest. 2016;126(2):653–66.

18. Oostindie SC, Van Der Horst HJ, Kil LP, Strumane K, Overdijk MB, Van Den Brink EN, et al. DuoHexaBody-CD37 ®, a novel biparatopic CD37 antibody with enhanced Fcmediated hexamerization as a potential therapy for B-cell malignancies. Blood Cancer J. 2020;10:30.

19. Deckert J, Park PU, Chicklas S, Yi Y, Li M, Lai KC, et al. A novel anti-CD37 antibody-drug conjugate with multiple anti-tumor mechanisms for the treatment of B-cell malignancies. Blood. 2013;122(20):3500–10.

20. Beckwith KA, Frissora FW, Stefanovski MR, Towns WH, Cheney C, Mo X, et al. The CD37-targeted antibody-drug conjugate IMGN529 is highly active against human CLL and in a novel CD37 transgenic murine leukemia model. Leukemia. 2014; 144(2):133.

21. Köksal H, Dillard P, Josefsson SE, Maggadottir SM, Pollmann S, Fane A, et al. Preclinical development of CD37CAR T-cell therapy for treatment of B-cell lymphoma. Blood Adv. 2019;3(8):1230–43.

22. Imai K, Takeuchi Y, Terakura S, Okuno S, Adachi Y, Osaki M, et al. Dual CAR-T Cells Targeting CD19 and CD37 Are Effective in Target Antigen Loss B-cell Tumor Models. Mol Cancer Ther. 2024;23(3):381–93.

23. Caulier B, Joaquina S, Gelebart P, Dowling TH, Kaveh F, Thomas M, et al. CD37 is a safe chimeric antigen receptor target to treat acute myeloid leukemia. Cell Reports Med. 2024;5(6).

24. Witkowska M, Smolewski P, Robak T. Investigational therapies targeting CD37 for the treatment of B-cell lymphoid malignancies. Expert Opin Investig Drugs. 2018;27(2):171–7.

25. Oostindie SC, Van Der Horst HJ, Kil LP, Strumane K, Overdijk MB, Van Den Brink EN, et al. DuoHexaBody-CD37 ®, a novel biparatopic CD37 antibody with enhanced Fc mediated hexamerization as a potential therapy for B-cell malignancies. Blood Cancer J. 2020;10:30.

26. van der Horst HJ, Oostindie SC, Cillessen SAGM, Gelderloos AT, Overdijk MB, Nijhof IS, et al. Potent Preclinical Efficacy of DuoHexaBody-CD37 in B-Cell Malignancies. HemaSphere. 2021;5(1):e504.

27. Lapalombella R, Yeh YY, Wang L, Ramanunni A, Rafiq S, Jha S, et al. Tetraspanin CD37 directly mediates transduction of survival and apoptotic signals. Cancer Cell. 2012;21(5):694–708.

28. Fujimura T, Yamashita-Kashima Y, Kawasaki N, Yoshiura S, Harada N, Yoshimura Y. Obinutuzumab in combination with chemotherapy enhances direct cell death in cd20positive obinutuzumab-resistant non-hodgkin lymphoma cells. Mol Cancer Ther. 2021;20(6):1133–41.

29. Mone AP, Cheney C, Banks AL, Tridandapani S, Mehter N, Guster S, et al. Alemtuzumab induces caspase-independent cell death in human chronic lymphocytic leukemia cells through a lipid raft-dependent mechanism. Leukemia. 2006;20:272–9.

30. Zhao X, Lapalombella R, Joshi T, Cheney C, Gowda A, Hayden-Ledbetter MS, et al. Targeting CD37-positive lymphoid malignancies with a novel engineered small modular immunopharmaceutical. Blood 2007;110(7):2569–77.

31. Ivanov A, Beers SA, Walshe CA, Honeychurch J, Alduaij W, Cox KL, et al. Monoclonal antibodies directed to CD20 and HLA-DR can elicit homotypic adhesion followed by lysosome-mediated cell death in human lymphoma and leukemia cells. Blood. 2009;114(20):4157–41.

32. Beers SA, French RR, Chan HTC, Lim SH, Jarrett TC, Vidal RM, et al. Antigenic modulation limits the efficacy of anti-CD20 antibodies: Implications for antibody selection. Blood. 2010;115(25):5191–201.

33. Lim SH, Vaughan AT, Ashton-Key M, Williams EL, Dixon S V., Chan HTC, et al. Fc gamma receptor IIb on target B cells promotes rituximab internalization and reduces clinical efficacy. Blood. 2011;118(9):2530–40.

34. Whittaker KC, Huang RP. Antibody Arrays Methods and Protocols. 2021. 45–54 p. http://www.springer.com/series/7651.

35. Pal Singh S, Dammeijer F, Hendriks RW. Role of Bruton’s tyrosine kinase in B cells and malignancies. Mol Cancer. 2018;17(1):1–23.

36. Benoit A, Bou-Petit E, Chou H, Lu M, Guilbert C, Luo VM, et al. Mutated RASassociating proteins and ERK activation in relapse/refractory diffuse large B cell lymphoma. Sci Rep. 2022;12(1).

37. Uddin S, Hussain AR, Siraj AK, Manogaran PS, Al-Jomah NA, Moorji A, et al. Role of phosphatidylinositol 3′-kinase/AKT pathway in diffuse large B-cell lymphoma survival. Blood. 2006;108(13):4178–86.

38. Höpken UE, Rehm A. Targeting the Tumor Microenvironment of Leukemia and Lymphoma. Trends in Cancer. 2019;5(6):351–64.

39. Wang Y, Wang C, Cai X, Mou C, Cui X, Zhang Y, et al. IL-21 Stimulates the expression and activation of cell cycle regulators and promotes cell proliferation in EBV-positive diffuse large B cell lymphoma. Sci Rep. 2020;10(1):1–13.

40. Jones ELL, Demaria MCC, Wright MDD. Tetraspanins in cellular immunity. Biochem Soc Trans. 2011;39(2):506–11.

41. Wood B, Sikdar S, Choi SJ, Virk S, Alhejaily A, Baetz T, et al. Leukemia & Lymphoma Abundant expression of interleukin-21 receptor in follicular lymphoma cells is associated with more aggressive disease Abundant expression of interleukin-21 receptor in follicular lymphoma cells is associated with more aggressive disease. Leuk Lymphoma. 2013;54(6):1212–20.

42. Zöller M. Tetraspanins: Push and pull in suppressing and promoting metastasis. Nat Rev Cancer. 2009;9(1):40–55.

43. Kumari S, Devi G V, Badana anil, ramesh Dasari V, rao malla rama. CD151-A Striking Marker for Cancer Therapy. Biomark Cancer. 2015;(7):7–7.

44. Vences-Catalán F, Kuo CC, Rajapaksa R, Duault C, Andor N, Czerwinski DK, et al. CD81 is a novel immunotherapeutic target for B cell lymphoma. J Exp Med. 2019;jem.20190186.

45. Yoshimura T, Miyoshi H, Shimono J, Nakashima K, Takeuchi M, Yanagida E, et al. CD37 expression in follicular lymphoma. Ann Hematol. 2022;101(5):1067–75.

46. Roumenina LT, Daugan M V., Noe R, Petitprez F, Vano YA, Sanchez-Salas R, et al. Tumor cells hijack macrophage-produced complement C1q to promote tumor growth. Cancer Immunol Res. 2019;7(7):1091–105.

47. Nemenoff RA, Markiewski MM. The role of complement in the tumor microenvironment Fac Rev. 2021;10(80).

48. Luo S, Wang M, Wang H, Hu D, Zipfel PF, Hu Y. How Does Complement Affect Hematological Malignancies: From Basic Mechanisms to Clinical Application. Front Immunol. 2020;11.

49. Song G, Cho WC, Gu L, He B, Pan Y, Wang S. Increased CD59 protein expression is associated with the outcome of patients with diffuse large B-cell lymphoma treated with R-CHOP. Med Oncol. 2014;31(7).

50. Fowler NH, Cheah CY, Gascoyne RD, Gribben J, Neelapu SS, Ghia P, et al. Role of the tumor microenvironment in mature B-cell lymphoid malignancies. Haematologica. 2016;101(5):531–40.

51. Singhal S, Rao AS, Stadanlick J, Bruns K, Sullivan NT, Bermudez A, et al. Human Tumor–Associated Macrophages and Neutrophils Regulate Antitumor Antibody Efficacy through Lethal and Sublethal Trogocytosis. Cancer Res. 2024;84(7):1029–47.

52. Bouwstra R, He Y, De Boer J, Kooistra H, Cendrowicz E, Fehrmann RSN, et al. CD47 expression defines efficacy of rituximab with CHOP in non–germinal center B-cell (Non-GCB) diffuse large B-cell lymphoma patients (DLBCL), but not in GCB DLBCL. Cancer Immunol Res. 2019 ;7(10):1663–71.

53. Zhao X, Lapalombella R, Joshi T, Cheney C, Gowda A, Hayden-Ledbetter MS, et al. Targeting CD37-positive lymphoid malignancies with a novel engineered small modular immunopharmaceutical. Blood. 2007;110(7):2569–77.

54. Rafiq S, Siadak A, Butchar JP, Cheney C, Lozanski G, Jacob NK, et al. Glycovariant antiCD37 monospecific protein therapeutic exhibits enhanced effector cell-mediated cytotoxicity against chronic and acute B cell malignancies. MAbs. 2013;5(5):723–35.

55. Wang J, Xu-Monette ZY, Jabbar KJ, Shen Q, Manyam GC, Tzankov A, et al. AKT Hyperactivation and the Potential of AKT-Targeted Therapy in Diffuse Large B-Cell Lymphoma. Am J Pathol 2017;187:1700–16.

56. Arribas A, Napoli S, Munz N, Stussi G, Bertoni F. PI3Kδ activation, IL6 over-expression, and CD37 loss cause resistance to naratuximab emtansine in lymphomas. Blood Adv. 2024; bloodadvances.2023012291.

57. Krause G, Baki I, Kerwien S, Knödgen E, Neumann L, Göckeritz E, et al. Cytotoxicity of the CD37 antibody BI 836826 against chronic lymphocytic leukaemia cells in combination with chemotherapeutic agents or PI3K inhibitors. Br J Haematol. 2016;173(5):791–4.

58. Rødland GE, Melhus K, Generalov R, Gilani S, Bertoni F, Dahle J, et al. The Dual Cell Cycle Kinase Inhibitor JNJ-7706621 Reverses Resistance to CD37-Targeted Radioimmunotherapy in Activated B Cell Like Diffuse Large B Cell Lymphoma Cell Lines. Front Oncol. 2019;9:1301.

59. Wilson WH, Young RM, Schmitz R, Yang Y, Pittaluga S, Wright G, et al. Targeting B cell receptor signaling with ibrutinib in diffuse large B cell lymphoma. Nat Med. 2015;21(8).

60. Wilson WH, Wright GW, Huang DW, Hodkinson B, Balasubramanian S, Fan Y, et al. Effect of ibrutinib with R-CHOP chemotherapy in genetic subtypes of DLBCL. Cancer Cell. 2021;39(12):1643–1653.e3.

61. Oostindie SC, Taylor RP, Van Der Horst HJ, Lindorfer MA, Cook EM, Tupitza JC, et al. CD20 and CD37 antibodies synergize to activate complement by Fc-mediated clustering. Haematologica. 2019 ;104(9):1841.

